# Identifying Robust Subclonal Structures through Tumor Progression Tree Alignment

**DOI:** 10.64898/2026.02.25.708046

**Authors:** Jacob Gilbert, Chih Hao Wu, Marina Knittel, Alejandro A. Schäffer, Salem Malikić, S. Cenk Sahinalp

**Affiliations:** Department of Computer Science, University of Maryland, College Park, MD, USA; Cancer Data Science Laboratory, National Cancer Institute, National Institutes of Health, Bethesda, MD, USA; Department of Computer Science and Engineering, UC San Diego, San Diego, CA, USA

## Abstract

Understanding and comparing tumor evolutionary histories is fundamental to cancer genomics. Clonal trees, used to model tumor progression, are rooted, unordered trees in which each node represents a subclone labeled by a set of distinct mutations.

To compare two clonal trees, we introduce `omlta`, the <u>o</u>ptimal <u>m</u>ulti-label tree <u>a</u>lignment, which removes the minimum number of mutation labels from the trees, so that the remaining trees are isomorphic. Computing `omlta` is NP-hard. Here, we present an algorithm to compute the `omlta`, with a running time of 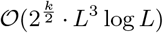 where *L* ≥ 1 is the total number of mutation labels occurring in the input trees and *k* is the minimum possible number of mutation labels that need to be removed for the alignment.

Our implementation (https://github.com/algo-cancer/omlta) is the first computational tool for determining the *optimal* alignment between clonal trees. We applied `omlta` to 126 cases from the TRACERx study on non-small cell lung cancers and some melanoma single-cell data.

## 1 Introduction

A sequence - an ordered list of labels (characters/symbols) - is a fundamental data type in computational biology; in fact, aligning two sequences was among the very first computational problems tackled in biology (1; 2; 3). At its core, the *sequence alignment* problem asks to compute the maximum number of matching labels which appear in the same sequential order in both input sequences. In this form, sequence alignment corresponds to the longest common subsequence of the two input sequences. More generalized forms of alignment may give different weights to different aligned label pairs including mismatched labels.

The dual problem to sequence alignment is the *edit distance*, which can be defined as the minimum number of label deletions from the input sequences so that the remaining sequences are identical. This definition is equivalent to the more commonly used definition of the edit distance as the minimum number of label insertions and deletions to be applied to a sequence to transform it to another sequence. Another related measure, the Levenshtein edit distance, differs slightly by permitting label substitutions in addition to insertions and deletions (1). It is well known that one can compute the optimal alignment of two sequences in time quadratic with their length (2). Any algorithm that computes the sequence alignment, implicitly computes the edit distance as well.

Trees, another data type commonly used in computational biology, generalize sequences by allowing branching, e.g., to model evolutionary history. One can generalize sequence alignment to alignment of rooted, node-labeled unordered (i.e., sibling nodes in the trees have no left-to-right ordering), trees as follows. Given two such trees, the *tree alignment problem* seeks the maximum-size subset of labeled nodes that induce a pair of isomorphic trees after all nodes and labels not in the output tree alignment set are deleted from the two input trees.

In this work, we are interested in aligning the evolutionary histories of tumor cells, which are commonly represented by *clonal trees* of tumor progression. In a clonal tree, each node corresponds to a genetically distinct population of cells, referred to as a (sub)clone. Formally, a clonal tree is a node-labeled tree where the sibling nodes are unordered, (ii) each node can be labeled with 0, 1, or more mutations, and (iii) each mutation label appears once in the tree. The labels of a given node represents the set of mutations which were acquired for the first time by its corresponding (sub)clone - without homoplasy. This follows from the observation that independent acquisition of an existing mutation in the tumor by another cell is rare (4). For some algorithms on clonal trees, it is necessary to assume that the root has no mutation labels as it may correspond to a set of normal cells that predated any somatic alterations, but we do not make such an assumption.

### Our main contributions

We introduce `omlta`, the *optimal multi-label tree alignment* for a pair of clonal trees (as well as any other tree type where a node can have any number of labels, provided that each label in the tree is unique) as the alignment which deletes the minimum possible number of labels so that the resulting trees are isomorphic (subject to empty node removals and node expansions; see Section 4.1 for the formal definition). Because `omlta` identifies structural features (e.g., subtrees) shared between input trees, it can be employed for making robust biological inferences through the use of clonal trees inferred via different methods on the same input data, or using different data obtained from the same sample, or using different samples from the same tumor.

We provide the first algorithm to compute the `omlta` between two clonal trees *T*_1_ and *T*_2_ and an implementation. The running time of our algorithm is 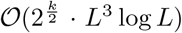, where *L* is the total number of distinct mutation labels in *T*_1_ and *T*_2_, and *k* is the value of the *optimal multi-label tree edit distance* (`omltd`) between *T*_1_ and *T*_2_, i.e., the number of mutation labels deleted by `omlta`. We can assume that *L* ≥ 1, so that log *L* is defined, because there would be no reason to construct a clonal tree for a tumor with no mutations. In our primary application, the trees *T*_1_ and *T*_2_ necessarily have the same mutation and label set, but that will not be the case for all uses of our method. Therefore, we point out that any mutation label occurring in exactly one of the two trees must be deleted as part of the alignment and hence `omltd`(*T*_1_, *T*_2_) must be at least as large as the symmetric difference of the two sets of mutation labels. This running time is polynomial when *k* is bounded by a constant and hence we show that the `omlta` problem is fixed parameter tractable (FPT). Since we also show that computing `omlta` (and `omltd`) is NP-hard, an FPT algorithm such as ours is qualitatively the best one can hope for with respect to efficiency. (See Section 4 and the Supplementary Information for the algorithm and some running time analysis.) The exponential term, 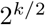, in the running time of our algorithm is strictly better than the 2.62^*k*^ term in the running time 𝒪 (poly(*n*) · 2.62^*k*^) of the state-of-the-art unordered tree edit distance algorithm for general (i.e., including, but not necessarily unique labeled) trees (5) where *n* is the number of nodes.

We applied `omlta` to clonal trees inferred from both multi-sample bulk and single cell sequencing data. On a cohort of 126 metastatic non-small cell lung cancer tumors from the TRACERx project (6), our algorithm could compute the `omlta` of clonal trees inferred by `CONIPHER` (7) (the method employed by the TRACERx project) and `PairTree` (8) (another popular method for clonal tree inference) on the same tumor quickly, often within seconds. We used `omlta` to assess the concordance between the trees inferred by these two methods, which we observed to vary substantially across the two major subtypes of NSCLC, namely LUAD (LUng ADenocarcinoma) and LUSC (LUng Squamous Cell carcinoma). Specifically, `omltd` values tend to be larger for the LUAD cases than the LUSC cases, implying that the trees inferred for LUAD cases are less reliable for downstream interpretation. Importantly we observed that `omltd` is negatively associated with the mean cancer cell fraction (CCF) among mutations in a tumor - implying that trees for tumors with lower CCF tend to be less robust to the choice of tree inference method. (b) Our algorithm could also efficiently compute the `omlta` of the clonal trees for the B2905 preclinical melanoma cell line (9) inferred (i) on data obtained by two distinct single cell sequencing technologies, or on data from the cell line before and after it was subjected to immunotherapy, or (iii) using different tree inference methods. As can be expected, the trees obtained from single cell data turned out to be more robust than those obtained from bulk data, even when the mutations were identified through transcriptome sequencing (see Section 2).

We also demonstrated the robustness of `omlta`*/*`omltd` compared to alternative similarity measures (see Sections A.6 and A.7). Our implementation is available at https://github.com/algo-cancer/omlta.

### 1.1 Related Works

#### Alignment Trees vs Consensus Trees

While we provide the first clonal tree alignment algorithm, `omlta` is related to frequently studied consensus trees (10; 11; 12; 13; 14; 15). Given a set of input trees, the consensus tree is the tree that minimizes the sum of distances to each input tree. A key difference between consensus tree algorithms and tree alignments is that typically, in an effort to make runtimes tractable, the considered distance for a consensus tree often counts only parent-child or ancestor-descendant relationships as in (10; 11; 15) whereas alignment trees are derived from the dual problem of `omltd` (see Section A.7 for a comparison of `omltd` to other distance measures). A further difference is that consensus trees must include *all* shared mutations across all input trees. As a consequence, given two input trees, *T*_1_ and *T*_2_, that are isomorphic except for the placement of a single mutation *a* - located on node *v*_1_ in *T*_1_ and on its descendant *v*_2_ in *T*_2_ - the consensus tree may collapse all nodes between *v*_1_ and *v*_2_ into a single node, thereby losing the ancestor–descendant relationships among the associated mutations that *T*_1_ and *T*_2_ otherwise share (14). In contrast, the alignment tree contains precisely those mutations that agree in ancestor-descendant relationship and topological location such that the induced trees are isomorphic; e.g., the `omlta` of *T*_1_ and *T*_2_ above excludes *a* but keeps all remaining mutations and their relative ordering.

#### Alternative Measures of Clonal Tree Similarity/Distance

For clonal tree similarity, the closest measure to `omltd` in the prior literature is *multi-label tree dissimilarity* (MLTD) proposed in (16). However, MLTD allows only leaf label deletions. One feature of MLTD is an edit operation denoted *node expansion*, which replaces a chosen node by a linear chain of two or more nodes and distributes its mutation labels among the nodes in the chain. Both MLTD and `omltd` permit the node expansion operation free of cost since trees of tumor progression inferred through the use of different sequencing technologies and/or methods may have varying levels of granularity/coverage. For example, while trees inferred from bulk sequencing data usually contain fewer numbers of nodes with multiple mutations per node (17; 18), trees inferred from single-cell data can be as detailed as a fully resolved clonal tree, also known as a *mutation tree*, containing only one mutation label per node (19; 20). By allowing node expansions at no cost, MLTD (and `omltd`) compute the similarity between clonal trees while reconciling such differences between them without penalty.The algorithms for computing MLTD and `omlta` can be modified to charge a penalty for this operation; they can also be modified to forbid node expansions, without any increase in asymptotic runtime.

In (16), the general tree edit distance was noted to be NP-hard, and MLTD was offered as a restricted but practical variant, solvable in polynomial time. Later in (20), a different version of the tree edit distance, between a clonal tree (representing an inferred phylogeny) and a mutation tree (where each node is assigned a single unique label) was considered and reduced to the general tree edit distance problem. Our algorithm for `omltd` is the first efficient solution to the unordered tree edit distance problem for two clonal trees allowing general node deletions and node expansions.

There are other measures of similarity based on the relative placement of labels, especially for species trees, including the well-known Robinson-Foulds distance (21). Because in species trees only the leaves are labeled - each with a single label representing a species, the measures defined to compare species trees are not commonly used to compare clonal trees (22). We include one such distance in our later comparisons, namely the Bourque distance (23), which is a generalization of Robinson-Foulds for clonal trees. Other approaches for comparing clonal trees (22; 24) are not based on a single measure but use a complementary pair of measures (e.g., ancestor-descendant and different-lineage measures). Furthermore, they lack the concreteness of the similarity evidence provided by the tree alignment itself. See Section A.6 for a detailed description of these similarity measures and Section A.7 for their comparison with `omltd` on simulated data.

#### Available Algorithms for Tree Edit Distance

The dual problems of tree alignment and tree edit distance, first proposed by (25), have been studied in many contexts. For rooted, *ordered* trees, several polynomial-time approaches have been developed (26; 27; 28; 29), and the state-of-the-art is a sub-cubic polynomial time algorithm (30). Computing the alignment and edit distance between *unordered* trees (such as clonal trees) however is NP-hard (31). Several works in the literature have proposed various workarounds to avoid intractable computation times for unordered tree edit distance (5; 32; 33; 34; 35). On the theoretical side, some families of trees which admit polynomial-time solutions have been identified and several fixed-parameter tractable algorithms have been proposed. Akutsu et al. (5) provided an 𝒪 (poly(*n*)· 2.62^*k*^)-time dynamic programming solution for unordered tree edit distance where the *k* is the actual tree edit distance of the input trees and *n* is the size of the input trees. Additionally, they showed that there is a polynomial-time solution when the maximum degree of the input trees is constant. The runtime and setting of (5) is most similar to our own; we provide an in-depth discussion of it and other theoretical unordered tree edit distance literature in Section A.1.

#### Alignment Tree vs Maximum Agreement Subtree

The `omltd` problem we study is also related to a well-studied problem called the *Maximum Agreement Subtree* problem (36; 37; 38). To our knowledge, this problem has been rarely studied in the context of cancer (39), but it has several non-cancer biological applications in its own right. We discuss in Section A.2 why the tree alignment and maximum agreement subtree problems are distinct and why algorithms for maximum agreement subtree do not transfer to our setting.

## 2 Results

We present applications of `omlta` to multi-sample bulk sequenced (bWES) lung cancer and single-cell sequenced (SCS) melanoma data sets. Additional results on simulated data (Section A.7), runtime (Section A.8), and a comparison of `omltd` to other measures (Sections A.6 and A.7) are available in Supplementary Materials.

### 2.1 Application to bWES trees: TRACERx non-small cell lung cancer cohort

#### Dataset and experimental setup

Al Bakir et. al presented clonal trees inferred for a cohort of n=126 metastatic non-small cell lung cancers (NSCLC) from the TRACERx project (6). The cohort includes both lung adenocarcinoma (LUAD) and lung squamous cell carcinoma (LUSC) cases (Fig. 3a). For each case, multiple spatially distinct samples from both primary and metastatic sites were subjected to bulk whole exome sequencing (bWES). The mutations in each case were clustered using `PyClone` (40) with some additional pre-processing (e.g., for removing “biologically improbable” clusters) and a clonal tree was inferred on these clusters using `CONIPHER` (7). Briefly, `CONIPHER` executes topological search through first performing subgraph pruning, then exhaustive enumeration and scoring of remaining trees towards finding an optimal-scoring solution. Several alternative models, objectives, and inference methods can, however, yield different tree reconstructions. Previous work (7) attempted to explore this variability by comparing the `CONIPHER` inferred tree for a single case from the NSCLC cohort to those inferred by three alternative methods, namely `PairTree` (8), `CITUP` (17), and `LICHeE` (41), and assessing the level of their similarity. However, since no method for tree alignment was available at that time, the study did not identify where the trees agreed exactly.

Here, we use `omlta` on *all* 126 metastatic NSCLC cases to align each tree obtained by `CONIPHER` to those by `PairTree` - the most recently developed method among the three alternative approaches employed in the original study, which was also reported to differ from `CONIPHER` the most (7). On simulated data, it was reported that `CONIPHER` agreed best with `CITUP` - with `LICHeE` being a close second (7). These two tools are now more than 10 years old and `CONIPHER` was developed to address their limitations, especially with respect to scalability: e.g., the number of mutation clusters that could be handled by `CITUP` is restricted to 10 and many of the cases in the TRACERx cohort have larger number of clusters.

We provide an in-depth exploration of the concordance between these two methods and the robustness of the clonal trees reported by the TRACERx NSCLC study (6). `PairTree` is methodologically distinct from `CONIPHER` as it performs sampling to search through the space of trees for maximizing the posterior probability of mutation pairs with ancestor-descendant relationship. For each case in the cohort, the input we provided to `PairTree` was exactly the same set of mutational clusters used for `CONIPHER`. Then we used `PairTree` to obtain the top 10 trees with the highest reported likelihood, and aligned each tree with that obtained by `CONIPHER` via `omlta`, to identify the tree with the lowest “discordance”, defined as the `omltd` normalized by the number of mutations observed in the tumor, for further consideration. Our choice to compare the `PairTree` tree that had the smallest `omltd` to the corresponding tree inferred by `CONIPHER` was to give the most favorable outcome to both tools; even in comparing the most similar trees between these methods, they still differ substantially.

See Fig. 1 for an example tumor with the top 10 trees obtained by `PairTree`, their pairwise discordances, as well as their discordances with the `CONIPHER` tree. As shown, the 10 trees inferred by `PairTree` differ not only from the tree obtained by `CONIPHER` but also from one another. These 10 trees appear to form two distinct clusters, within which the discordance among trees is lower than that between clusters: {3, 4, 5, 7, 8, 10} and {1, 2, 6, 9 }. The latter cluster also exhibits lower discordance with the `CONIPHER` tree.

**Figure 1.**
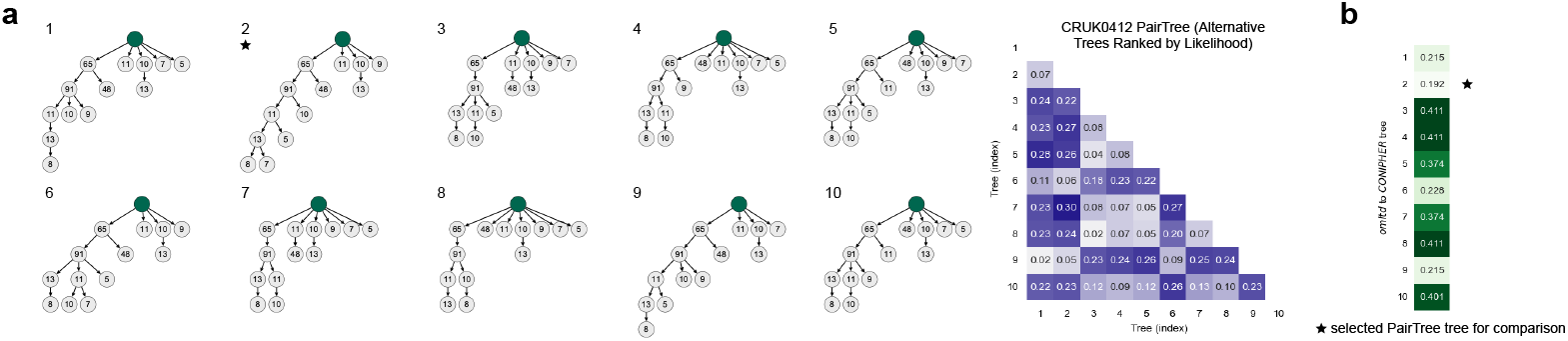
For each of the 126 metastatic NSCLC cases in the TRACERx cohort (6), we used `PairTree` to obtain an alternative tree to that reported by `CONIPHER` on the same set of mutational clusters employed by `CONIPHER` in the original study. **a.** First, for each case, all `PairTree` produced trees were evaluated and the top 10 trees according to their reported likelihood were identified: the left panel depicts the top 10 trees for case CRUK0412 (out of 1795 trees sampled by `PairTree`) - on each node, the number of mutations in the cluster assigned to that node is provided. The right panel depicts the discordance between each pair of trees - defined as the `omltd` normalized by the total number of mutation labels. **b**. Next, each of the top 10 `PairTree` trees for each case is aligned with the `CONIPHER` tree, and the one with the lowest discordance is identified. The panel shows for case CRUK0412, the discordance between each `PairTree` tree and the `CONIPHER` tree. Since, with respect to its reported likelihood, tree 2 has the lowest discordance, it is identified as the `PairTree` alternative for this case.

### Inferred topological structures and their biological implications vary among alternative tools

`CONIPHER` and `PairTree` inferred trees vastly differ in their overall topology (Section A.4). Using the average node depth of a tree *T* (more precisely, 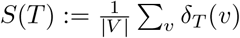 - where δ (*v*) is the depth of a node *v*) as a measure of “tree imbalance” (42), we found that `CONIPHER` trees are significantly more imbalanced than the corresponding `PairTree` trees (Wilcoxon signed-rank test, *P* = 2.2 × 10^−22^; Fig. 2b). This is likely due to `CONIPHER`’s subgraph pruning approach, which strongly preserves the nested structure of mutational clusters and thereby favors imbalanced trees. In contrast, `PairTree`’s sampling-based approach imposes less stringent hierarchical constraints.

**Figure 2:**
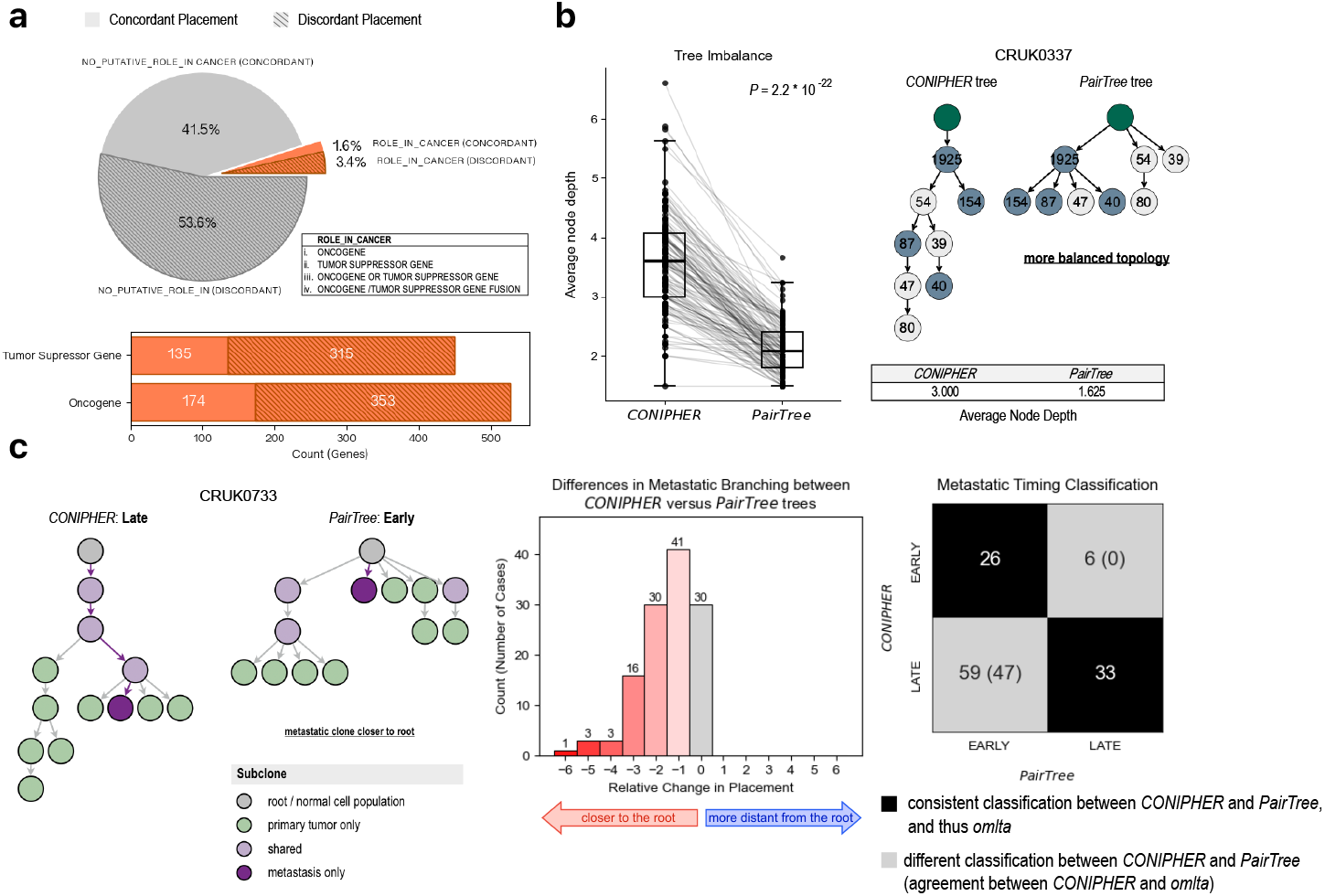
**a**. Distribution of mutations across the metastatic NSCLC cases from the TRACERx cohort. *Top panel* : a mutation is either in a gene with a “putative role in cancer” (i.e., is a oncogene or tumor suppressor - colored orange) or with “no putative role in cancer” (colored gray). Orthogonally, the placement of the mutation is either concordant (i.e., is preserved in the `omlta` of the trees inferred by `CONIPHER` and `PairTree` for the case - solid color) or discordant (i.e., is deleted in the `omlta` - shaded color). The proportion of discordant mutations are higher among those in genes with a role in cancer. *Bottom panel* : the proportion of concordantly placed mutations among tumor suppressor genes are similar to that among oncogenes. **b**. For any given case, the “imbalance” of a clonal tree, measured as the mean node depth (distance from the root), is typically higher if inferred by `CONIPHER`. E.g., for case CRUK0337, the imbalance of the tree inferred by `CONIPHER` is 3, but the imbalance of the tree inferred by `PairTree` is 1.625. (Blue nodes in the trees represent the mutational clusters preserved in their `omlta`.) **c**. The inferred timing of metastatic branching (6) differs between the `CONIPHER` and `PairTree` trees for case CRUK0733. *Left panel* : In the `CONIPHER` inferred tree, the metastatic branching is characterized as “late” whereas in the `PairTree` tree for the same case, it is characterized as “early”. *Middle panel* : The timing of metastatic branching differs significantly between `CONIPHER` and `PairTree` trees: `PairTree` typically places the metastatic branching closer to the root. *Right panel* : The number of cases for which `CONIPHER` and `PairTree` agree/disagree with respect to the timing of metastatic branching; additionally, the timing of the metastatic branching based on `omlta` trees (given in parentheses) agree with `CONIPHER` more often.

Topological differences can substantially affect biological interpretation. For example, analysis of the structure of subclones in a cancer may guide combination therapy (43) and may yield insights about changes in the rate and types of mutations in later (lower in the tree) subclones (44; 45). For these data, the TRACERx NSCLC study (46) classified each metastatic branching (i.e., seeding) event as either “early” or “late” based on whether mutations “clonal” across primary tumor samples were respectively absent from (early) or shared with (late) the corresponding metastatic sample. Hong and colleagues first showed that different methods of tumor phylogenetic inference on the same input data can yield different interpretations about metastasis in a patient (47). While the study did not differentiate mutations with respect to their clonality beyond whether they are present (or not) in all primary samples, one can use the inferred clonal tree to categorize all mutations that are on the trunk of the tree, i.e., prior to any branching within the primary tumor, as “clonal”. This allows a more refined temporal characterization of metastatic branching events (as early vs. late clonal) based on the node on the trunk of the tree for the primary tumor from which the metastatic sample branches out. (See, e.g., Fig. 2c, left panel). In trees inferred by `PairTree`, metastatic branching typically occurs closer to the root compared to trees inferred by `CONIPHER` (Fig. 2c, middle panel). There are 59 cases where metastatic branching would be characterized as ‘late’ using `CONIPHER` trees but would be characterized as ‘early’ using `PairTree` trees. A more robust characterization is possible via the alignment trees: `omlta` trees characterize 47 of these 59 cases as late (in agreement with `CONIPHER`) and characterize the remaining 12 as early (in agreement with `PairTree`). Interestingly, an additional six cases characterized as ‘early’ by `CONIPHER` are characterized as ‘late’ by `PairTree`; the `omlta` classification agrees with `PairTree` in all six. There are two remaining cases which were not characterized as ‘early’ or ‘late’ by by `PairTree` because their trees had no metastasis specific branch from the trunk of the tree. (Fig. 2c, right panel; see Supplementary Table S1 for details.)

#### Robustness of mutational placement in clonal trees vary across tumors

We investigated mutational placements that are robust to the choice of inference method by aligning trees obtained from the same case by `CONIPHER` and `PairTree`. Pairwise alignment is a label-preserving bijection between subsets of vertices of two labeled trees that induces isomorphic subtrees on the set of preserved labels while minimizing the number of omitted labels. For clonal trees, preserved and omitted mutational labels respectively correspond to ‘concordant’ and ‘discordant’ mutational placements.

We used `omlta` to quantify the robustness of mutational placements between trees inferred by `CONIPHER` and `PairTree` on each tumor from the TRACERx dataset (6; 46). When stratified by histological subtype of NSCLC, the trees obtained by `CONIPHER` and `PairTree` exhibited significantly higher discordance for LUAD cases than LUSC cases (Mann-Whitney U test, *P* = 0.003; Fig. 3b and 3c). To identify the basis of this difference, we first considered the cancer cell fraction (CCF) of mutations across the cases of each subtype. We observed that LUAD cases typically harbor a larger fraction of low CCF (typically subclonal) mutations than LUSC cases; e.g., in several LUAD cases more than half of all mutations had CCF ≤ 0.3 (See Supplementary Fig. S5, top panel). This was consistent with the findings of the TRACERx study (46): “in LUAD, the majority of frequently mutated cancer genes were subject to significant positive subclonal selection … no significant positive selection was detectable in truncal mutations.”

**Figure 3:**
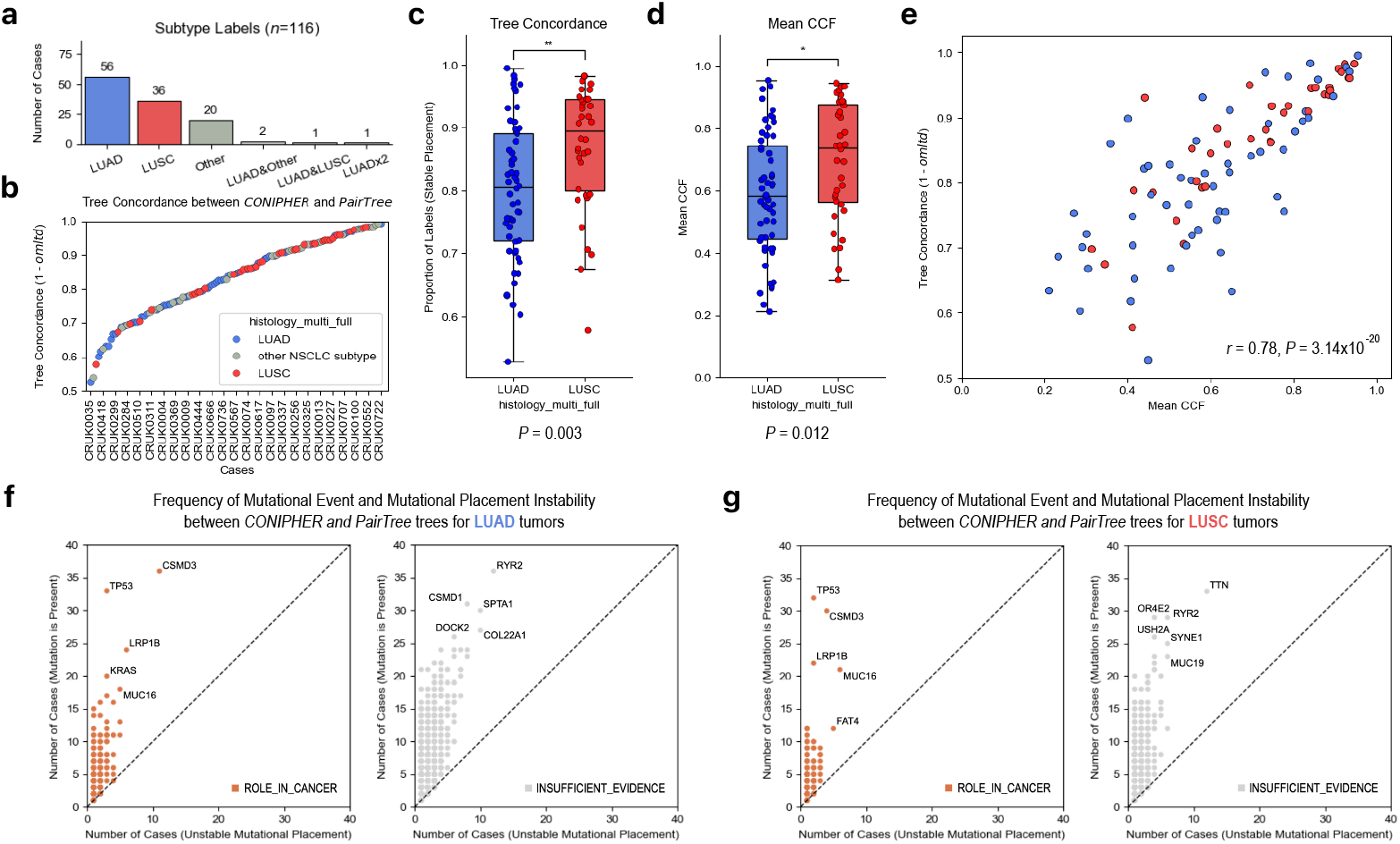
**a**. Metastatic tumors from the TRACERx NSCLC cohort, consists of 56 LUAD cases and 36 LUSC cases; the cohort also includes an additional 24 cases of other subtypes and 10 cases with no subtype annotation. **b**. For each case, `omlta` between the trees inferred by `CONIPHER` and `PairTree` was obtained and their concordance was measured as the proportion of robust mutational placements, i.e., 1 - `omltd`/number of mutations. The panel depicts all cases with a subtype annotation (only a subset of cases are labeled explicitly due to limited space) sorted based on the concordance values: LUSC cases tend to have higher concordance values and thus are more robust. **c-d**. A comparison between the distribution of LUAD and LUSC cases with respect to **c** concordance values, and **d** mean CCF of the mutations harbored: both distributions reveal a significant difference between the two subtypes. **e**. Pearson correlation between the mean CCF and tree concordance values among LUAD and LUSC cases: higher mean CCF values among LUSC cases may explain why their inferred trees tend to have higher concordance. **f-g**. For the two NSCLC subtypes, **f** LUAD and **g** LUSC, scatterplots show the number of cases in which a gene is mutated (y-axis) against the number of cases in which a mutation within that gene has discordant placement between `CONIPHER` and `PairTree` trees (x-axis). Genes with known “role in cancer” (in orange) are shown separately from genes with no putative role (in gray). The points for outlier genes are labeled to the extent that labels could be added without overlaps.

Next, we explored whether topological differences between the trees inferred by either method, as well as the `omlta` trees can explain the differences between LUAD and LUSC cases. For any given case, the tree inferred by `CONIPHER` was typically more imbalanced than the tree inferred by `PairTree` - with their `omlta` falling in between. However within the trees inferred by each method (`CONIPHER`, `PairTree` or their `omlta`), the imbalance distribution for LUAD cases was similar to that for LUSC cases. (See Supplementary Fig. S6.) Likewise, the imbalance difference between the `CONIPHER` tree and the `PairTree` tree is distributed similarly among LUAD cases and LUSC cases.

The concordance between the trees inferred by `CONIPHER` and `PairTree` for a given case is strongly associated with the subtype (Mann-Whitney U test, *P* = 0.003; Fig. 3c); however, this association can be attributed to LUAD cases harboring a higher proportion of low CCF mutations (Mann-Whitney U test, *P* = 0.012; Fig. 3d), which are typically subclonal. In fact, there is a strong association between mean CCF and tree concordance (Pearson’s Correlation coefficient = 0.78, *P* = 3.14 × 10^−20^; Fig. 3e). While it is possible to infer the temporal ordering of clonal mutations robustly, the placement of a subclonal mutation may differ among inference methods, especially if they are based on different assumptions and parameter settings. As a result, cases with higher mean CCF appear to be more robust to the choice of tree inference method.

#### Robustness of mutational placement in clonal trees vary among genes with distinct roles in cancer

Next, we evaluated the robustness of mutational placements according to the known roles of the genes harboring them in the context of cancer. The genes that harbor these mutations are annotated using a combination of two cancer gene censuses, Catalogue Of Somatic Mutations In Cancer (COSMIC) (48) and OncoKB (49; 50). Below, we refer to mutations by their genes and do not distinguish different mutations in a case - primarily for compactness of display. If a gene harbors more than one mutation in a case, we regard the gene as having unstable placement if any of its mutations were deleted from the `omlta`.

An earlier study from the TRACERx consortium showed that driver mutations among such consensus cancer genes are typically clonal - with exceptions (51). On the other hand the `CONIPHER` tool used in the TRACERx NSCLC study does not utilize annotations of genes harboring the mutations in their placement. Thus, we perform a post-hoc analysis of the placement of mutations harbored by genes with a role in cancer.

Genes previously characterized as an oncogene and/or tumor suppressor gene, or as a gene fusion are collectively referred to as genes having a ‘role in cancer’ and are distinguished from those with ‘no putative role’ in tumor initiation, proliferation, or metastasis. In aggregate, 56.9% of genes (3.4% ‘role in cancer’ and 53.5% ‘no putative role’) had discordant inferred placements in at least one case in the NSCLC cohort. It is expected that genes which implicated in tumor progression are associated with clonal expansion—implying that all topological orderings that include such mutations should consistently assign them to more clonal nodes in a tree, and thereby be robust across multiple solutions. Yet such genes exhibited discordant mutational placement between trees in our comparisons with a 1.65-fold higher rate than other genes. Specifically, genes with a ‘role in cancer’ had disproportionately more discordant placements between trees (discordant:concordant ratio of 2.13:1; 3.4% vs 1.6%) compared to genes without an established functional role in cancer (discordant:concordant ratio of 1.29:1; 53.5% vs 41.5%). There was not a discernible difference in the relative number of discordantly placed genes between oncogenes and tumor suppressor genes (Fig. 2a).

Subsequently, we examine whether frequently mutated genes have robust inferred mutational placements. As can be expected, the number of cases in the TRACERx cohort in which a mutated gene is discordantly placed by `PairTree` and `CONIPHER` (i.e., the x-axes in Fig. 3f,g) increases with the number of cases in which that gene is mutated (i.e., the y-axes in Fig. 3f,g). Furthermore, commonly mutated genes with a role in cancer (colored orange) are as likely (about one in four) to be discordantly placed as other commonly mutated genes (colored gray). A subset of these genes likely represent key mutations in lung cancer. Interestingly, *TP53* which is commonly mutated in NSCLC cases is not often placed discordantly (it was placed discordantly in only 6/89 cases) possibly because it is typically mutated clonally.

A key purpose of inferring clonal trees is to aid oncologists in designing combination therapies such that each major subclone is treated. Among the commonly mutated genes labeled in Fig. 3f,g, somatic mutations in several genes have been associated with response to different treatments. For example, presence of a mutation in either *MUC16* (52) or *LRP1B* (53; 54) is directly associated with better response to immune checkpoint blockade. Such associations with treatment response make correct placement of subclonal mutations essential to suggest how immune checkpoint inhibitors should be combined with targeted therapies to target subclonal variations in disjoint subtrees (43). Several of the genes labeled in Fig. 3f,g have been shown to be associated with higher tumor mutational burden (TMB) including *LRP1B, TP53, CSMD3, MUC16, RYR2, TTN*, but not *KRAS* (55; 56). The associations with high TMB for *RYR2* and *TTN*, which are not among the cancer related genes in COSMIC or OncoKB, serve as a reminder that these curated sources have very strict rules for which genes are classified as having a role in cancer, but including other genes in tumor phylogenetics analysis has value.

The quest to identify subtrees that have gained a “mutator phenotype” (57) with a particular high rate of some types of mutations, such as microsatellite instability (58; 59), was one of the motivating problems for the field of tumor phylogenetics at its inception at the end of the 20th century (59). Therefore, the association with high TMB for certain mutated genes above is important to test assumptions about which subtrees have low or high mutation rates. The association with high TMB is also important clinically since higher TMB also predicts response to immune checkpoint inhibitors, as indicated by the pan-cancer FDA approval of immune checkpoint blockade to treat any cancer that has TMB greater than 10/Mbp. Correctly placing in clonal trees the (mutated) genes associated with higher TMB is also useful because the subtrees with these mutations at the root likely have a distinct, higher mutation rate (44). For these gene-specific and general reasons, the discrepant placement of the labeled genes such as *MUC16*, as compared to *TP53*, limits what can be robustly inferred from the clonal trees for some patients in the TRACERx NSCLC dataset. For those patients and samples for which `CONIPHER` and `PairTree` agree on the placement of these frequent mutations, more robust inferences about tumor progression can be made.

### 2.2 Application to SCS trees: B2905 preclinical melanoma model

We applied `omlta` to tumor progression trees inferred from three sequencing datasets of tumors derived from the B2905 cell line, a murine model of UV-induced, RAS-mutated human melanoma (9): (i) a bulk whole exome sequencing (bWES) dataset of 24 single cell derived sublines, (ii) a bulk whole transcriptome sequencing (bWTS) dataset of the same 24 sublines, and (iii) three single-cell whole transcriptome (scWTS) datasets described in (14). We discuss the `omlta` between trees derived from these datasets below.

#### Aligning trees for 24 single cell derived sublines individually sequenced using bulk methods

Our first set of experiments are on bulk sequencing datasets. As described in detail below, we (i) compare a pair of trees, each reconstructed using a distinct tree inference method applied to the the same bulk sequencing dataset, and also (ii) compare a pair of trees, each reconstructed by the same tree inference method, applied independently to two distinct datasets, which were both obtained from the same tumor.

The bWES dataset consists of 24 single-cell-derived sublines, each sequenced independently, featuring a total of 1689 mutations. Each of the 24 sublines was derived from an isolated single cell from the parental cell line and grown *in vitro*, and represents a genetically homogeneous cell population. As such, the 24 sublines represent an idealized, high coverage version of whole exome sequencing data from 24 single cells. Formally, the input genotype matrix representing the bWES dataset consists of 24 rows, each representing a single subline, and 1689 columns, each representing a mutation observed in one or more of the sublines. Each entry of the binary matrix indicates the absence or presence of a mutation in a subline.

We applied to the bWES dataset two computational methods for inferring trees from single-cell sequencing data: `ScisTree` (60) (see Fig. 4a for the tree inferred by `ScisTree`) and `SCITE` (19) (see Fig. 4b for the tree inferred by `SCITE`). While these methods are methodologically similar, they exhibit some key differences in their tree-scoring strategies and in the approaches they employ for exploring tree space. Neither `ScisTree` nor `SCITE` provides a theoretical guarantee of optimality within a specified time.

**Figure 4.**
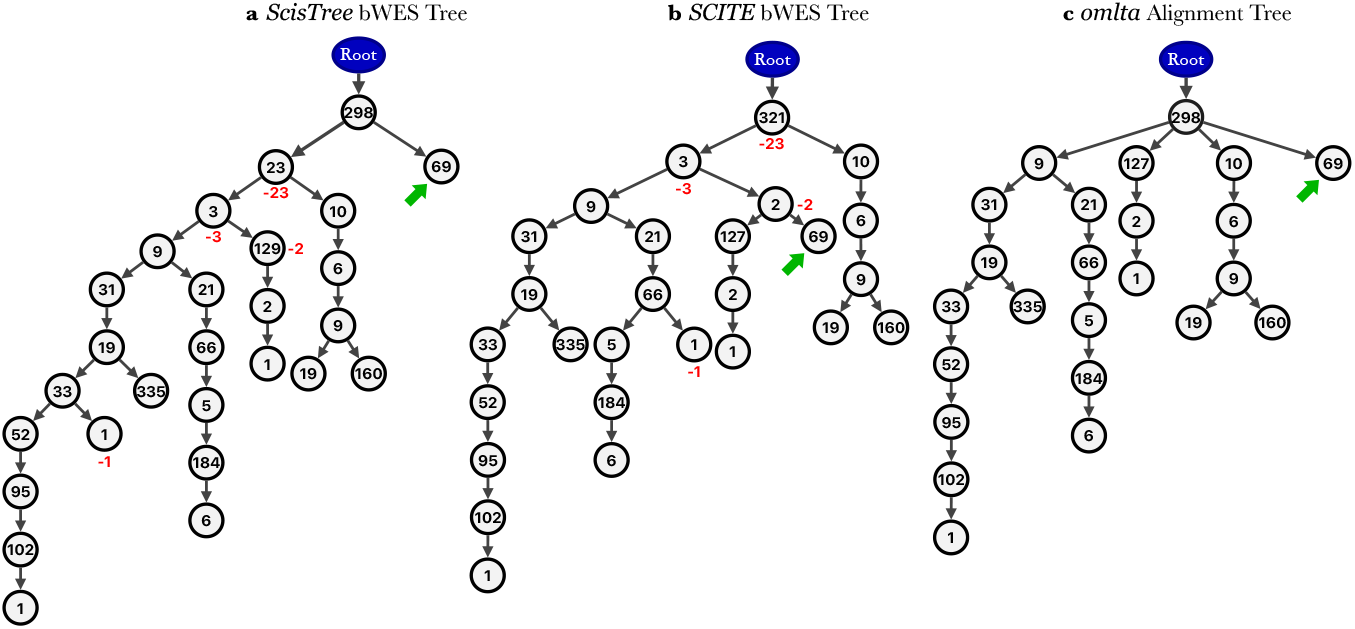
Comparison of trees reconstructed from bWES data consisting of 24 sublines and 1689 mutations by the use of **a** `ScisTree` (60) and **b** `SCITE` (19). Each non-root node (clone) is labeled by the number of mutations occurring for the first time at that node. When aligning these trees, `omlta` removes 29 mutation labels. Numbers shown in red next to the nodes indicate the number of mutations deleted from those nodes. **c.** The `omlta` tree between the trees shown in **a** and **b**. The green arrows point to the same mutational cluster across all trees.

The alignment between the trees inferred by `ScisTree` and `SCITE` on the same input dataset demonstrate high topological concordance in terms of mutational ordering, with only a few mutation label deletions (29 out of 1689). The main difference between the two trees is in the placement of matching nodes each with (the same set of) 69 mutations. In the tree shown in Fig. 4a, the 69 mutations are labels in the right child of the root, while in the tree shown in Fig. 4b, the 69 mutations are in the root’s left child’s subtree; these are indicated by green arrows). To maximize concordance, `omlta` removes those mutations that are on the path from the root to the node with these 69 mutations in the tree shown in Fig. 4b - although these mutations are concordantly placed in the two trees–enabling the retention of the larger set of 69 mutations in the alignment tree.

Next, we compared trees reconstructed using the same method on distinct sequencing datasets, each derived from the same parental cancer cell line. We conducted two experiments, differing in the sets of mutations used for tree inference, as explained below. The first sequencing dataset used in these experiments is the bWTS dataset, consisting of 11 sublines in which 536 mutations were detected. Each of these 11 sublines was also sequenced using bulk whole-exome sequencing; i.e., these 11 sublines are included in the bWES dataset described above. To allow for a direct comparison, we restricted the bWES dataset to the 11 sublines common to both bWTS and bWES datasets, obtaining the *reduced-bWES* dataset, consisting of 11 sublines and 1420 mutations.

In principle, the bWTS and reduced-bWES datasets differ technologically in sampling depth and sequencing noise, which may potentially result in discrepancies in the inferred ancestral relationships between mutation pairs across their respective trees. To demonstrate that our `omlta` algorithm can effectively reconcile trees derived from sequencing data collected by different wet lab methods, we conducted two experiments. The first experiment was on trees inferred using the restricted set of mutations common to both the BWTS and the reduced-bWES datasets. The second experiment was on trees inferred using the mutations observed in either dataset.

The set of mutations observed in both bWTS and reduced-bWES datasets contains only 92 mutations. Due to this reduced size, we were able to use a tree inference method with optimality guarantee, namely `PhISCS` (20), to build the trees. The `omltd` between the bWTS (Fig. 5a) and reduced-bWES (Fig. 5b) trees (both on 11 sublines and 92 common mutations) had a notably low value of 12, due to 6 mutation label deletions in each tree. The `omlta` tree (Fig. 5c) features the maximal set of concordant mutations across bWTS and reduced-bWES trees. If a pair of mutations are not ordered in one input tree, due to being labels of the same node, while the mutations appear in different nodes in the other input tree, then the order implied by the latter input tree is featured in the output alignment tree. As such, `omlta` can resolve the temporal ordering of each retained mutation pair assigned to the same node in exactly one input tree by using the mutation ordering in the other input tree, potentially producing an output tree with higher resolution than either input tree.

**Figure 5.**
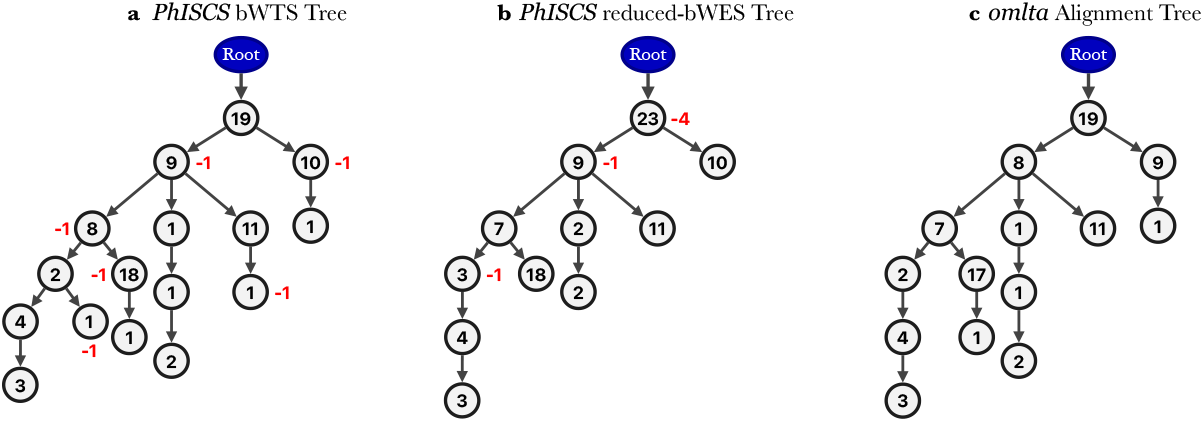
Comparison of clonal trees reconstructed by `PhISCS` (20) from (***a***) bWTS and (***b***) reduced-bWES data, each with 11 sublines, featuring a total of 92 mutations common to both datasets. Each non-root node (clone) is labeled by the number of mutations occurring for the first time at that node. When aligning these trees, `omltd` reports 6 mutation label deletions - numbers shown in red next to the nodes indicate the number of mutations deleted from those nodes. (***c***) The `omlta` alignment tree between the trees shown in *a* and *b*.

As a companion to the analysis of bulk sequencing melanoma trees built by `PhISCS`, we built trees with the complete set of mutations observed in either bWTS (11 sublines, 536 mutations) and reduced-bWES (11 sublines, 1420 mutations) datasets. Since tree inference methods with an optimality guarantee, such as `PhISCS`, can not handle 1420 mutations, we employed `ScisTree` for tree inference. The `omlta` alignment tree between the bWTS tree (Figure 6a) and the reduced-bWES tree (Figure 6b) is shown in Figure 6c. Focusing on the 92 mutations shared between the bWTS and reduced-bWES trees (as the other mutations are necessarily excluded from the alignment tree), we observe that only 6 of these 92 mutations are not included in the alignment tree–concordant with the alignment between the bWTS and reduced-bWES trees inferred on mutations common to both datasets (see Fig. 5).

**Figure 6.**
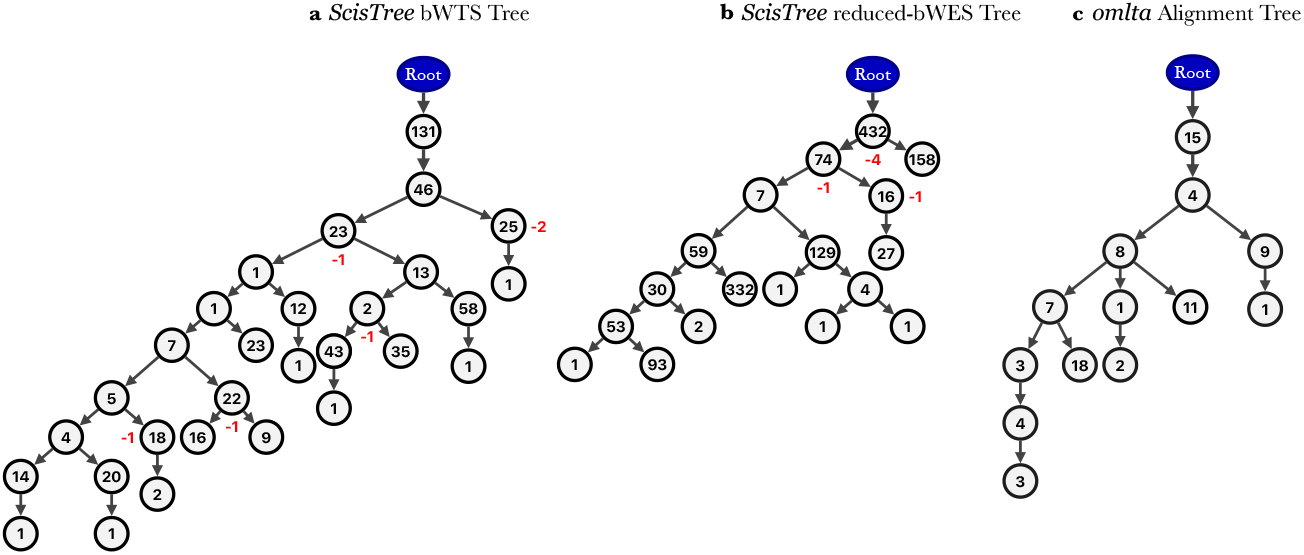
Comparison of clonal trees reconstructed by `ScisTree` (60) on **a** bWTS (11 sublines and 536 mutations) and **b** reduced-bWES (11 sublines and 1420 mutations) data. Each non-root node (clone) is labeled by the number of mutations occurring for the first time at that node. In addition to the mutations present in only one of the two trees, when aligning these trees, `omlta` also reports 6 mutation label deletions in each of them - numbers shown in red next to the nodes indicate the number of mutations deleted from those nodes. **c.** The `omlta` alignment tree between the trees shown in **a** and **b**.

The comparisons of trees built with `PhISCS` and `ScisTree` demonstrate that the tree reconstructed from bulk transcriptome data closely resembles that of the tree inferred from the bulk exome data obtained from the same set of sublines. Notably, the `omltd` for `PhISCS`-built trees on mutations shared between the two datasets is equal to that for `ScisTree`-built trees on the complete set of mutations observed in each dataset. This demonstrates that `ScisTree`, a method with no optimality guarantees, produced results on a larger set of mutations that are comparable to those produced by an optimal tree reconstruction method, `PhISCS`, on a restricted set of mutations - providing a strong support for trees built by `ScisTree` on these datasets. In fact, the subtrees induced on the set of common mutation labels for the reduced-bWES trees built by `PhISCS` (Fig. 5b) and `ScisTree` (Fig. 6b) are isomorphic and the corresponding bWTS trees (Fig. 5a and Fig. 6a) have only two differently placed mutations (not shown).

#### Reconciling trees reconstructed from single-cell RNA sequencing datasets

We subsequently evaluated the trees built from single-cell datasets derived from the same parental cancer cell line. For that, we used two datasets from a preclinical study related to the impact of immunotherapy on intratumoral heterogeneity and subclonal evolution. Specifically, cells from the B2905 melanoma cell line were subcutaneously implanted into six syngeneic mice randomized to two groups corresponding to different treatment settings, mouse anti-CTLA-4 antibody treated (“Treated”, 4 mice) and mouse IgG2b as isotype, negative control (“Control”, 2 mice). Cells from each tumor were sequenced using the Smart-seq2 protocol (61) to generate single-cell, full-length transcript reads, resulting in our *smartseq-treated* and *smartseq-control* datasets comprising 508 and 163 cells, respectively. Application of `omlta` on these trees may offer insights into subclones and mutations they harbor which may have undergone immunoediting. In particular, if a node in the smartseq-control tree is missing from the smartseq-treated tree but its parent node is retained, then the subclone defined by that node may have been selectively eliminated by the immune system. Such a deletion points to a set of mutations which may induce neoantigens recognizable by the immune system.

We inferred clonal trees from these mutational profiles by the use of `ScisTree` (60)—for both the control and treated tumors; these trees are presented in Fig. 7a. Additional details related to clustering and parameters for tree inference are provided in Section A.5 of Supplementary Material. The topological differences between the true clonal trees for control and treated tumors would imply anti-CTLA-4 therapy induced immunoediting, as they correspond to the negative selection against subclonal mutations.

**Figure 7:**
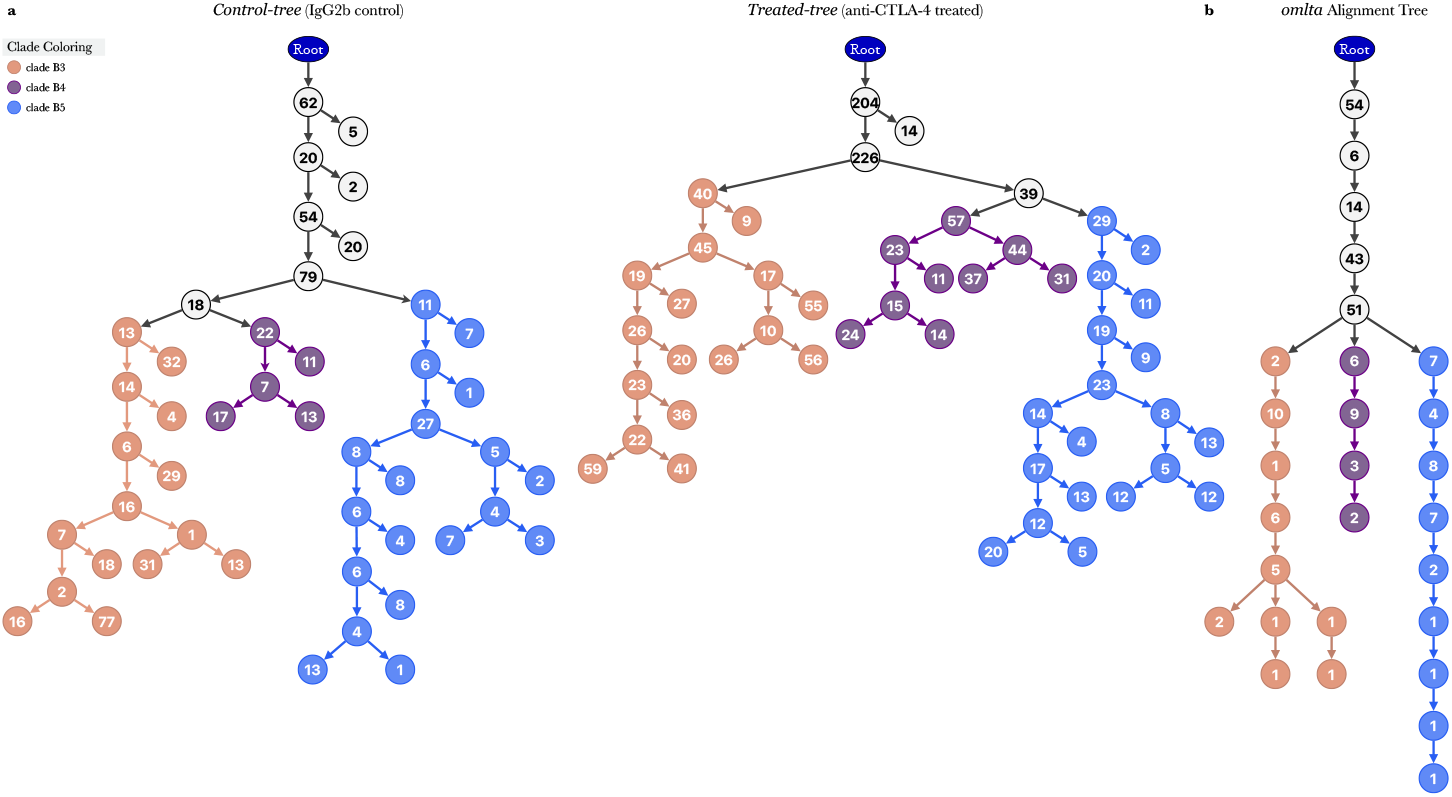
Tumor progression trees inferred by ScisTree on scRNA data from a preclinical study of anti-CTLA-4 immunotherapy for melanoma. a. Tumor progression trees inferred by `ScisTree` from *smartseq-control* data (“Control-tree”, left) and *smartseq-treated* data (“Treated-tree”, right). The clonal trees are labeled by the number of mutations that were inferred to have been acquired for the first time in the evolution of the tumor for each node/subclone. Based on shared mutations labeled at the root of each subtree, there are three major clades (B3, B4, and B5) that were previously identified in (14) and are colored accordingly. b. An optimal alignment between Control- and Treated-tree, “Alignment-tree”, as determined by computing `omltd`. The clonal tree is labeled by the number of common mutations with concordant placement between Control- and Treated-trees.

The resulting Control-tree was built on 24 clusters with 740 mutations, while the Treated-tree was built on 25 clusters with 1518 mutations; see Figure 7a. The number of mutations common to both trees was 583, among which 333 were deleted to obtain the `omlta`, i.e, the alignment tree obtained from these clonal trees featured 250 mutations, as presented in Figure 7b. Notably, the `omlta` exhibits a more “linear” topology than either clonal tree, primarily preserving a specific lineage from each of the three major subtrees (colored orange, purple and blue) observable in both the Control-tree and the Treated-tree. This indicates that, while the major clade structure is preserved between the Control-tree and the Treated-tree, the exact placements of mutations differ substantially. This is not surprising since clonal tree inference from these datasets is substantially more challenging than that from the bulk sequencing datasets not only because individual cells were sampled from highly heterogeneous tumors but also the read coverage on mutational loci was highly variable across cells.

In a final experiment, we compared tumor progression trees inferred exclusively from the control tumors, where each cell was sequenced using one of the two distinct single-cell RNA sequencing protocols: 163 single cells were sequenced using the Smart-seq2 protocol, forming our *smartseq-control* dataset. An additional 163 single cells were sequenced using the Seq-Well protocol (62) forming our *seqwell-control* dataset (see (14) for details). In Figure 8a, we present the trees we inferred for the two datasets, and in Figure 8b, we show their corresponding alignment tree. The `omltd` of these two trees and additional comparisons of trees reconstructed using different clustering criteria are provided in Tables 1 and 2.

**Figure 8:**
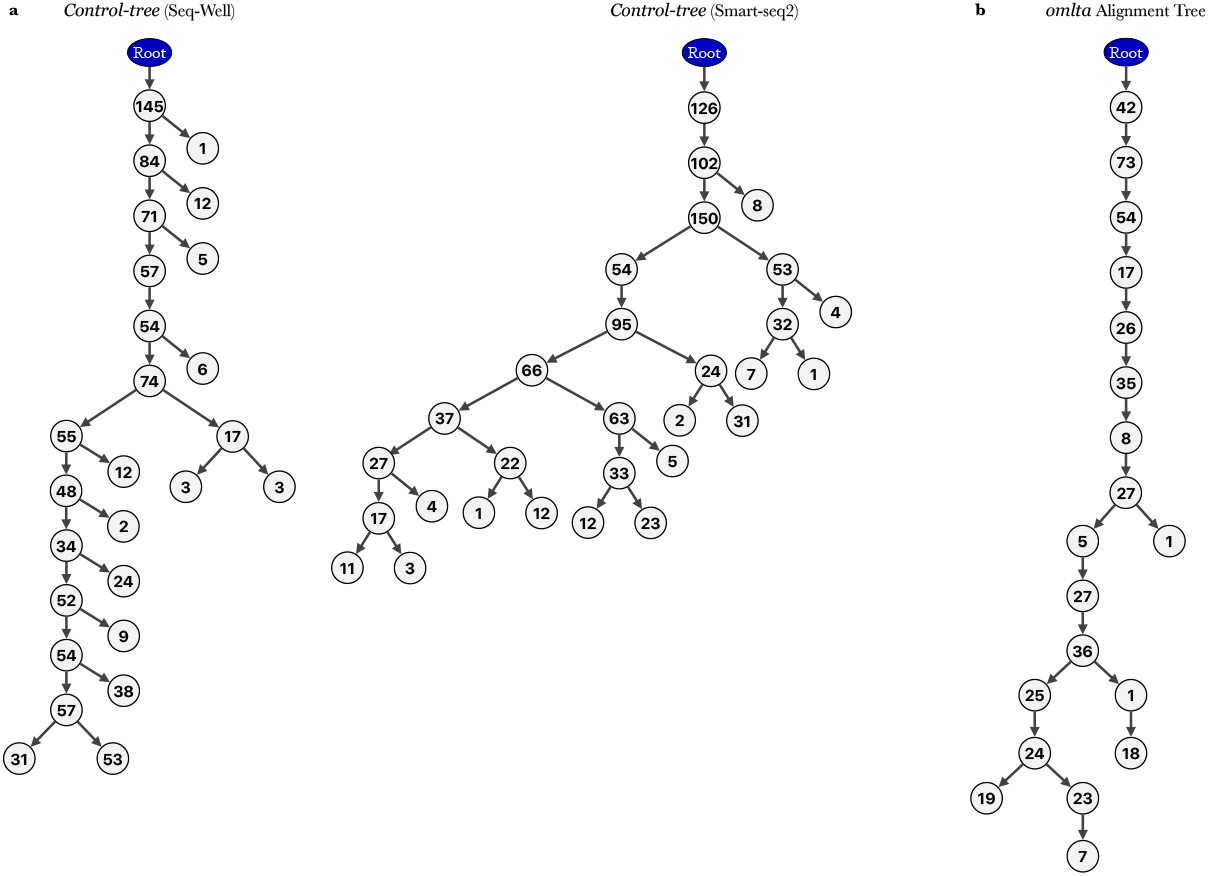
Tumor progression trees inferred by ScisTree on scRNA data from control tumors sequenced by Seq-Well and Smart-seq2. **a.** Tumor progression trees inferred by `ScisTree` from *seqwell-control* data (“Seq-Well”, left) and *smartseq-control* data (“Smart-seq2”, right). The clonal trees are labeled by the number of mutations that were inferred to have been acquired for the first time in the evolution of the tumor for each node/subclone. b. An optimal alignment between Seq-Well and Smart-seq2 tree, “Alignment tree”, as determined by performing `omlta`. The clonal tree is labeled by the number of common mutations with concordant placement between Seq-Well and Smart-seq2 trees.

**Table 1.** Pairwise comparisons of clonal trees reconstructed by the use of ScisTree from control and treated tumors sequenced by Smart-seq2. The `omltd`/2 for each pair of trees is provided; specifically, this is the number of labels that are deleted from one of the two trees.

| Clusters ($S=n$) | Min. cells per cluster | Control | | Treated | | Common Mutations | omltd/2 |
| --- | --- | --- | --- | --- | --- | --- | --- |
|  |  | mutations | clusters | mutations | clusters |  |  |
| 100 | 15 | 490 | 3 | 995 | 3 | 361 | 196 |
| 100 | 14 | 554 | 4 | 1122 | 4 | 426 | 248 |
| 100 | 13 | 554 | 4 | 1122 | 4 | 426 | 248 |
| 100 | 12 | 628 | 6 | 1304 | 6 | 505 | 306 |
| 100 | 11 | 699 | 9 | 1376 | 9 | 551 | 335 |
| 100 | 10 | 705 | 10 | 1393 | 10 | 554 | 338 |
| 100 | 9 | 720 | 15 | 1448 | 15 | 569 | 326 |
| 100 | 8 | 740 | 24 | 1518 | 25 | 583 | 333 |
| 100 | 7 | 741 | 30 | 1531 | 33 | 584 | 355 |

**Table 2.** Pairwise comparisons of clonal trees reconstructed by the use of ScisTree from replicate control tumors sequenced by Smart-seq2 and Seq-Well protocols (min. cells per cluster is 1). The `omltd`/2 for each pair of trees is provided; specifically, this is the number of labels that are deleted from one of the two trees.

| Clusters ($S=n$) | Seq-Well | | Smart-seq2 | | Common Mutations | omltd/2 |
| --- | --- | --- | --- | --- | --- | --- |
|  | mutations | clusters | mutations | clusters |  |  |
| 5 | 978 | 4 | 1025 | 5 | 978 | 136 |
| 6 | 983 | 5 | 1025 | 6 | 983 | 78 |
| 7 | 983 | 6 | 1025 | 7 | 983 | 72 |
| 8 | 983 | 6 | 1025 | 8 | 983 | 119 |
| 9 | 984 | 7 | 1025 | 8 | 984 | 311 |
| 10 | 994 | 8 | 1025 | 9 | 994 | 182 |
| 11 | 994 | 8 | 1025 | 10 | 994 | 365 |
| 12 | 994 | 9 | 1025 | 10 | 994 | 389 |
| 13 | 994 | 9 | 1025 | 11 | 994 | 395 |
| 14 | 995 | 10 | 1025 | 10 | 995 | 395 |
| 15 | 995 | 10 | 1025 | 12 | 995 | 430 |
| 16 | 995 | 11 | 1025 | 12 | 995 | 476 |
| 17 | 1000 | 12 | 1025 | 13 | 1000 | 501 |
| 18 | 1001 | 13 | 1025 | 14 | 1001 | 523 |
| 19 | 1001 | 13 | 1025 | 14 | 1001 | 606 |
| 20 | 1001 | 14 | 1025 | 16 | 1001 | 533 |

A key difference between the trees obtained for single-cell sequencing data and those obtained for single-cell-derived subline bulk sequencing data is that `omltd` values for the former are much higher; up to two-thirds of the mutation labels shared between the aligned trees may need to be deleted. This is partially because of sampling variability. While the single-cell-derived sublines are fixed (with each subline being well-represented in its respective bulk sequencing dataset), each single-cell dataset is comprised of distinct cells possibly representing distinct cell populations with many distinct mutations. Data sparsity plays a role as well, with many mutational loci not covered by even a single read in certain cells. In addition, pre-clustering of cells could add noise to tree inference process.

Despite these challenges, the `omlta` alignment between the single-cell trees retains a substantial number of mutations, reflecting that single-cell trees can provide reliable information about the temporal ordering of mutations and the branching patterns leading to distinct cell lineages. The lineages preserved in the `omlta` tree can be robustly leveraged in downstream analyses, such as integrating mutation profiles with gene expression data in single cells to identify potential associations. Perhaps more importantly, these experiments demonstrate that our `omlta` algorithm remains practical and effective even in the presence of extensive mutation label deletions in large single-cell trees, highlighting its practical utility.

## 3 Discussion

Robust tumor phylogeny inference remains fundamentally constrained by the limitations of current sequencing technologies and analytic approaches. Regarding sequencing technology, multi-sample bulk data often lack the resolution needed to unambiguously place key mutations, while single-cell sequencing suffers from substantial sparsity and technical noise that impede confident reconstruction of clonal relationships. Regarding analytical methods, there are different software methods for inferring clonal trees and mutation trees. Moreover, different parameter settings within the same method can yield markedly different trees from the very same dataset.

One premise of this work is that the variation in tree inference methods can be viewed as an opportunity rather than as a limitation. If the trees produced by two tree inference methods can be robustly compared by a neutral third method, then one can identify the substructures shared by the two inferred trees. To address the need for a tree comparison method, we introduced the notion of optimal tree alignment, `omlta`, for clonal trees of tumor progression and developed an efficient algorithm for computing an optimal alignment. Our method for computing optimal alignments naturally produces the integer distance between the two trees, `omltd`, which is the sum of the numbers of node label deletions that must be applied in each tree to make the resulting derived trees isomorphic. The `omltd` is unique, but there may be multiple optimal alignments, as illustrated below in Figure 9.

**Figure 9:**
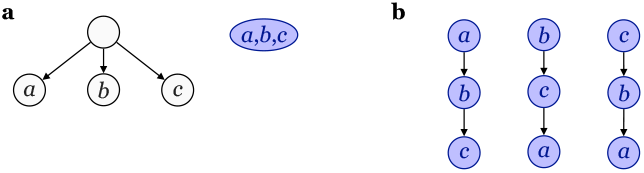
**a**. A star tree and a single node tree, which have an `omltd` distance of 4 between them. **b**. Some possible expansions of a single node tree with label set *a, b, c*.

Although we showed that computing `omlta` is formally intractable (NP-complete), we were able to compute `omlta` in practical runtimes for all input instances tested, including instances with more than 1000 mutations. Thus, the NP-completeness does not represent a practical limitation. However, one practical limitation of our methods and results is that `omlta` compares exactly two trees at a time. Since there are more than two methods for inferring clonal trees and these methods have parameter settings that can be varied to produce different clonal trees, it may be desirable to compare more than two trees simultaneously. Just as pairwise tree alignment corresponds to pairwise sequence alignment, the problem of comparing more than two trees corresponds to multiple sequence alignment, which is a widely studied problem with widely used methods that offer a mixture of formal optimization criteria and heuristics (63; 64; 65).

In conclusion, applying our framework to clonal trees inferred from both multi-sample bulk and single-cell sequencing data, we demonstrated utility of `omlta` in revealing robust evolutionary features and consistent mutational placements across pairs of methods and data types, thereby enabling more reliable downstream analyses. The use of `omlta` should lead to more rigorous use of clonal trees in cancer genomics. Our `omlta` framework will also provide an impetus to develop new methods for tree inference because it is now possible to compare more rigorously a new method to existing methods, as we compared `CONIPHER` to `PairTree`.

## 4 Methods

### 4.1 Preliminaries

In this paper, we consider rooted, unordered trees *T* = (*V, E*) with node set *V*, edge set *E*, and a specified root node denoted *r*(*T*). For two nodes *u* and *v, u* ≠ *v*, such that *u* lies on the path from *r* to *v*, we say that *u* is an *ancestor* of *v* and that *v* is a *descendant* of *u*. A node *v* ∈ *V* is a *child* of *u* ≈ *V* if *v* is a descendant of *u* and (*u, v*) ∈ *E*. For a node *u* ≈ *V*, `child`(*u*) denotes the set of children of *u*. For shorthand, *v* ∈ *T* is used with the same meaning as in *v* ∈ *V*, and |*T* | is used to denote the number of nodes in *T* (i.e., |*T* | = |*V* |). Given two nodes *u, v* ∈ *T, P* (*u, v*) denotes the set of nodes on the path from *u* to *v* in *T*, including *u* and *v*. We denote the subtree rooted at *v* ∈ *T* as *T* (*v*). A *forest* F is a collection of node-disjoint trees {*T*_1_, *T*_2_, …, *T*_ℓ_ }. For any *T*_*i*_ ≈ ℱ, ℱ \*T*_*i*_ denotes the forest ℱ excluding *T*_*i*_. The generalization of the problem to forests is needed because during our recursive algorithm single trees may be transformed into multiple trees via deletions of nodes and edges.

The nodes in our trees have labels from an alphabet ∑, which includes the mutations observed in the sample and to be assigned to nodes in the tree. Each node may have multiple labels (representing mutations), which are treated for computation purposes as an unordered set, but each label appears at most once in a tree. We let *λ* : *V* →2^∑^ represent a label function such that *λ*(*v*) is the label set of node *v*. Given a set of nodes *X*, we define *λ*(*X*) to be the set of all labels present in nodes in *X*, that is *λ*(*X*) =⋃ _*v*≈*X*_ *λ*(*v*). For a tree *T* = (*V, E*), we may refer to *λ*(*V*) as *λ*(*T*).

#### Definition 1.

*A* ***multi-label tree*** *T* = (*V, E, λ*) *with nodes V, edges E, and label function λ* : *V* → 2^∑^ *is an unordered tree in which for all v* ∈ *V*, |*λ*(*v*)| ≥ 1, *except the root which may have no label. Furthermore, for all v*_*i*_, *v*_*j*_ ≈ *V with i* ≠ *j, label sets λ*(*v*_*i*_) *and λ*(*v*_*j*_) *are disjoint. A multi-label forest* ℱ = {*T*_1_, *T*_2_, …, *T*_ℓ_} *is a collection of multi-label trees such that for any T*_*i*_, *T*_*j*_ ≈ ℱ *with i j, label sets λ*(*T*_*i*_) *and λ*(*T*_*j*_) *are disjoint*.

#### Definition 2.

*An* ***edit operation*** *for the multi-label tree edit distance problem is defined as one of the following tree operations:*

- ***Label Deletion***: *Delete a label a* ∈ *λ*(*v*) *for a node v* ∈ *V* . *We denote this label deletion by* `del`(*v, a*). *For a set of labels A* ⊆ *λ*(*v*), *let* `del`(*v, A*) *denote deletion from λ*(*v*) *of all labels that belong to A, i*.*e*., *if we start with B* = *λ*(*v*), *then after* `del`(*v, A*) *is applied, we have λ*(*v*) = *B* \ *A*.
- ***Node Deletion***: *Delete a node v* ≈ *V and, if v is not the root of T, make all nodes in* `child`(*v*) *children of u, where u denotes the parent of v. If v is the root of T, then all nodes w* ≈ `child`(*v*) *become roots of their own distinct trees T* (*w*) *and this transforms a single tree into a forest. The node deletion operation can be applied only to empty nodes. We denote this operation as* `del`(*v*).
- ***Expansion***: *For a node v with* |*λ*(*v*) | *>* 1 *and some A* ⊆ *λ*(*v*), *add a new node u to V, set λ*(*v*) ←*λ*(*v*) \*A and λ*(*u*) ←*A. If v was a non-root node with parent w, we set E* →*E*∪ {(*w, u*), (*u, v*)}\ (*w, v*). *Otherwise, u becomes the new root of T and we set E* ←*E*∪ {(*u, v*)} . *We denote such an expansion as* `expand`(*v, A*).

Given a forest ℱ with a node *v*, we use ℱ \ *v* to denote the forest ℱ after performing `del`(*v*). Similarly, given a forest ℱ and a subset *A* of labels present in ℱ, we use ℱ \*A* to denote the forest ℱ after performing `del`(*v*_*a*_, *a*) for all *a* ∈ *A*, where *v*_*a*_ is the node in ℱ such that *a*∈ *λ*(*v*_*a*_).

The edit distance between two trees is the cost of a minimum-cost sequence of edits, where an edit may be performed on either tree, that transforms both trees into the same tree, the alignment tree. With this in mind, we must assign costs to each edit. In this work, we are interested in finding the number (and the set) of labels that must be deleted in both trees; therefore, we set the cost of label deletion to 1 and the cost of empty-label node deletion and node expansion to 0, i.e., these are free operations. Since empty-label node deletion and node expansion are 0-cost operations, we say two forests are “equal” or “the same” when referring to two forests that could be transformed into equal forests after a sequence of only empty-label node deletions and node expansion. We make the distinction between forests that are exactly equal and forests that are equal after additional node expansions and empty-label node deletions when it is relevant to do so.

We can now formally define:

#### Problem 1.

*Given two multi-label forests ℱ* _1_, *ℱ* _2_, *the* optimal multi-label tree edit alignment (`omlta`*) asks to find a minimum-cost sequence of label deletions, node deletions, and expansions needed to transform* ℱ_1_ *and* ℱ_2_ *into equal forests, and* optimal multi-label tree edit distance (`omltd`*) of* ℱ_1_, ℱ_2_, *denoted as* `omltd`(ℱ_1_, ℱ_2_), *is the cost of this edit sequence*.

For the application of our `omlta` algorithm to real data, we are only interested in finding `omlta` for trees. However, we define the edit distance problem on forests because the intermediate states when edits are applied in sequence can include multi-tree forests and hence, forests are needed in the proof of correctness and in the formal analysis of asymptotic runtime. When given two trees as input, the minimum-cost edit sequence obtained by our algorithm, which we give in the following subsection, yields the same output tree (not a forest) derived by edits applied to one input tree or the other, and we refer to this common output tree as the *alignment tree*.

Before discussing the algorithm, we give a small insight into the limitations of the node expansion operation. It may seem that allowing node expansions at no cost may trivialize our distance computation and potentially allow any two trees to be matched without any label deletions needed. As a simple example contradicting this idea, consider two trees, a star and a single node, as in Figure 9a. Following the above definition, expansion can never create a node with degree greater than 1. Therefore, using expansion on a single node can only create various chain topologies with the mutation labels in various orders but can never create a star. Figure 9b shows some example expansions, and we may observe that one of the only notable changes in any expansion of the single node labeled *a, b, c* is the order of labels in the expanded trees. In all cases, the minimum number of labels that need to be deleted to transform the star and single node into the same tree is 4 (2 from each input tree). Essentially, `omltd` does not penalize long paths of nodes with degree 1 by allowing free node expansions, but does penalize actual topological differences in branching and mutation label ordering, which is precisely what we aim to measure with this distance.

### 4.2 An *O*(2^*L*/2^ · *L*^2^)-time recursive algorithm for omlta

In this section, we present a simplified recursive formula to help compute `omlta`*/*`omltd` in 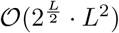-time where *L* is the number of mutations occurring in the two input trees (counting mutations in both trees as 1). The recursive formula identifies the minimum number of label deletions needed to induce two isomorphic trees up to node deletions and expansions. It is also constructed to handle forests as input since the recursive computation sometimes requires solving smaller subproblems in which the intermediate structure may be a forest of more than one tree. We denote the function to be computed as `omltd`(ℱ_1_, ℱ_2_) where _1, 2_ can be trees or forests. Given two trees *T*_1_, *T*_2_, `omltd`(*T*_1_, *T*_2_) finds the minimum number of label deletions that can be used to recover the alignment tree, solving both `omltd` and `omlta` simultaneously. After discussing runtime and correctness of our given recursive solution, we show in Supplementary Section 4.4 how to improve the exponent in the asymptotic running time to be linear in *k* rather than linear in *L*, where *k* = `omltd`(ℱ_1_, ℱ_2_). Our improved algorithm and runtime analysis require knowing an upper bound *B* = 𝒪 (*k*) on `omltd`(𝒪_1_, 𝒪 _2_) in advance, and so we run our algorithm with every possible value, 1, 2, 3, …, stopping when we reach *k*, and show that this search loop does not affect the runtime asymptotically.

Given two forests ℱ_1_ and ℱ_2_, the core idea of our algorithm is a recursive approach in which we iteratively target a single mutation label in the root of any tree of ℱ_1_ and find the node with its matching label in ℱ_2_ (if one exists). The algorithm then proceeds either to match or to delete this pair of nodes. In either case, the algorithm proceeds recursively on the remaining set of labels after removing the current considered label from both forests. If all the labels of a root node *v* of any tree in ℱ_1_ have been deleted by the algorithm or *v* is matched, then *v* is removed from ℱ_1_ for the remainder of the algorithm and the subtrees of the children of *v* are added into ℱ_1_ so that their labels may be matched or deleted next. By considering each label one at a time, all labels will eventually be matched or deleted while guaranteeing that the resulting forests after all deletions are equal as required by the formalized Problem 1. The algorithm finds exactly the sequence of matches and deletions that minimizes the number of label deletions needed to reach the alignment tree.

The main challenge is how to determine whether each label should be matched or deleted. For any tree *T* in ℱ_1_ that has no corresponding label-isomorphic tree in ℱ_2_, for each label in *r*(*T*), the algorithm will try both options, deleting and matching. Since there are *L* labels, we need to explore both options at most *L* times, which suggests a factor of 2^*L*^ in the runtime; actually, since the choice of whether to match or to delete arises only when the label occurs once in each forest, formal analysis in Section 4.3 shows that the exponential term can be reduced to 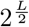. To improve the dependency further from *L* to *k*, we rely on the fact that the number of label deletions is bounded by some positive integer *k≥* `omltd`(ℱ_1_, *ℱ*_2_), and therefore, we argue that we only have to explore both choices at most *k* times, otherwise the distance must be larger than *k*.

Figure 11 shows an example of the basic approach of our algorithm. We compute `omltd` by iteratively matching or deleting each mutation label in the input trees. A root-to-leaf traversal of one of the input trees, chosen arbitrarily from the two, determines the order in which we consider mutation labels. In a preliminary step, we do a linear traversal of the trees and delete any mutation labels not present in both trees. In our example figure, there is one such label *m*_6_ only present in *T*_2_, which is deleted immediately. Then we begin our iterative approach to handle remaining mutation labels shared between the input trees. As mentioned, we start at the root of *T*_1_. First, the algorithm handles *m*_0_. Any mutation contained in the root of both trees may be preserved at no cost via a node expansion. Thus, *m*_0_ is matched between the two trees in the `omlta` and will remain in the final alignment tree. Next, *m*_1_, the other label in the root of *T*_1_ is considered. *m*_1_ is in a lower level of *T*_2_ and therefore matching *m*_1_ and retaining this label in the alignment tree requires some edits. The `omltd` computation therefore breaks into two alternative cases. In one alternative case, the algorithm tries to maintain *m*_1_ in both trees. As shown in the figure, maintaining *m*_1_ requires deleting the path of nodes above *m*_1_ from the current root of *T*_2_ and then also deleting the new hanging subtree of *m*_5_, since it will not have an isomorphic match in *T*_1_ if *m*_1_ is preserved. In the resulting trees, a node expansion can be made in each tree to place *m*_1_ in its own node to be retained in the resulting alignment tree. In the other alternative case, *m*_1_ is deleted in both trees. In either case, after handling *m*_1_, the algorithm recurses and continues iterating on the remaining trees. Whichever path leads to a lower number of total mutation label deletions is then chosen as the solution path and the alignment tree from that path is outputted. In the figure example, intuitively, deleting *m*_1_ is most likely the correct choice since it only requires 2 immediate deletions whereas matching *m*_1_ requires 86 immediate deletions.

Before giving our recurrence for computing `omlta`*/*`omltd`, we define a useful function that is used in the final case of the recurrence. Given a forest ℱ and node *v* in tree *T* ∈ ℱ, let *R*(ℱ, *v*) denote the forest ℱ after deleting all nodes of *P* (*v, r*(*T*)) (recall that *P* (*v, r*(*T*)) is the path in *T* from root *r*(*T*) to node *v*, including the endpoints) from ℱ and adding the subtree of any child node *c* ≈*/ P* (*r*(*t*), *v*) of a deleted node *p* ≈ *P* (*r*(*t*), *v*) \ {*v*}. See Figure 10 for an example.

**Figure 10.**
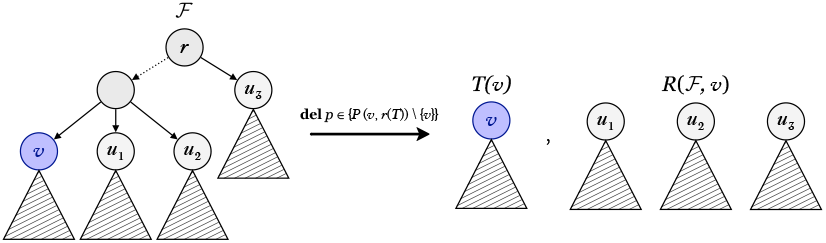
*Left panel:* A forest ℱ (in this example just a tree) with root *r* and node *v. Right panel:* Tree *T* (*v*) and forest *R*(ℱ, *v*) obtained from ℱ by deleting all nodes on the path from *r* to *v* and adding to ℱ the subtrees of any children node *c* ≈*/ P* (*r, v*) whose parent *p* ≈ *P* (*r, v*) *\* {*v*} has been deleted.

**Figure 11.**
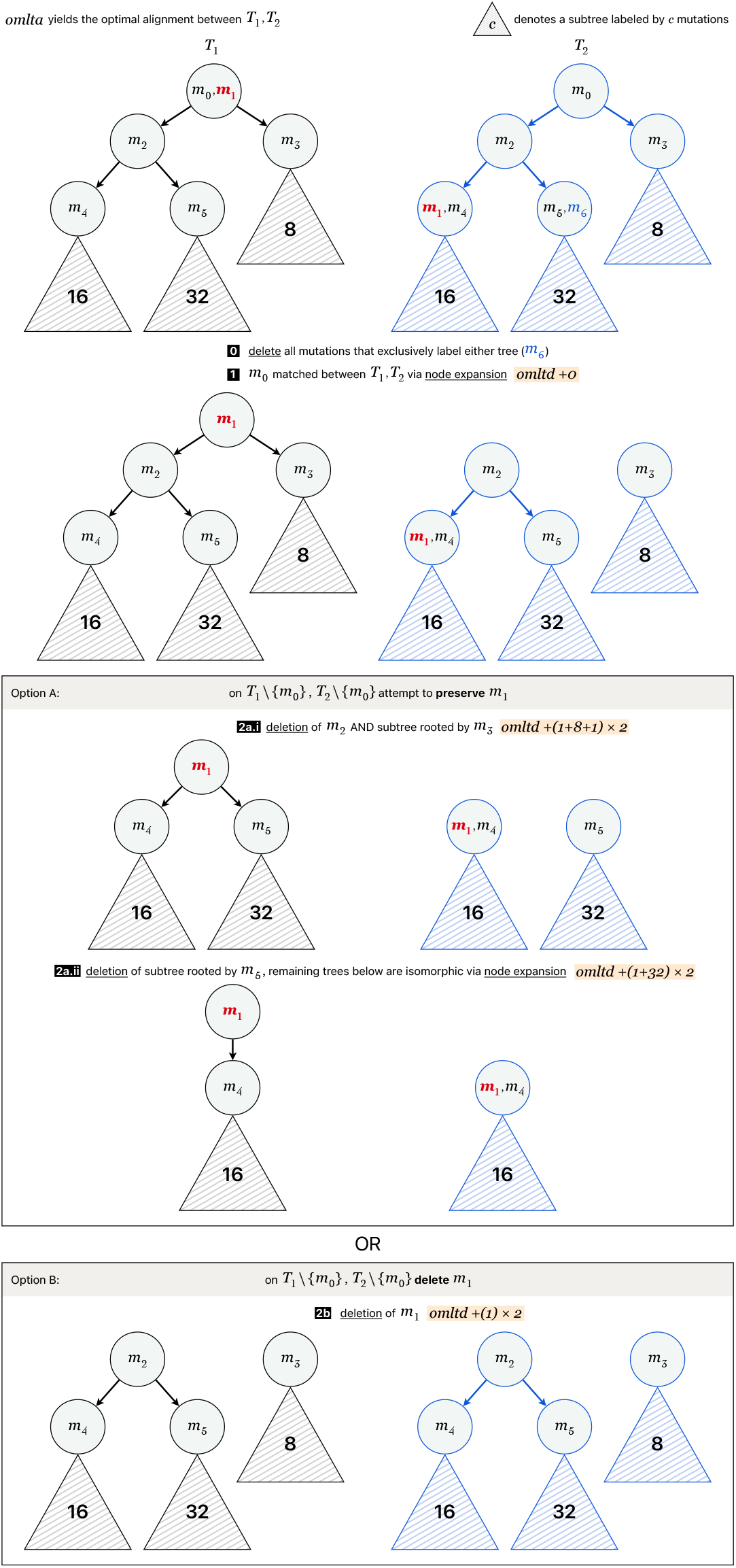

We consider one mutation at a time. Let *v*_1_ be the root of a tree *T*_1_ in ℱ_1_. Let *b* be a label in *v*_1_. Recall that empty-labeled node deletions have no cost towards `omltd`, so we may delete empty-labeled nodes for free. Additionally, if a root node of an input tree of ℱ_1_ (or ℱ_2_) has no labels, we may similarly delete that empty node before applying the recursive algorithm to ensure that all root nodes contain at least one label.

Let *u*_2_ be the node in tree *T*_2_ ∈ ℱ_2_ with *b* ∈ *λ*(*u*_2_) if it exists. If *u*_2_ exists, let *v*_2_ be the node in the path from *r*(*T*_2_) to *u*_2_, inclusive, that is closest to *r*(*T*_2_) and shares some label with *v*_1_, i.e., *λ*(*v*_1_) ∩ *λ*(*v*_2_) ≠ ∅. Set *a* to be any mutation in *λ*(*v*_1_) ∩ *λ*(*v*_2_) or just *b* if no matching label for *b* was found in ℱ_2_. While not written in formula 1, we assume that when the last label of a node is deleted at any step, the now empty-labeled node is deleted as well. We define the following recursive formula for `omltd`(ℱ_1_, ℱ_2_):

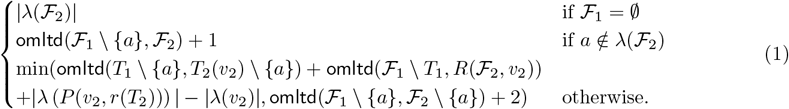

The above recurrence helps us compute `omltd` and find which labels need to be deleted for `omlta`. The three above cases handle all possibilities for label *a*. In the first case, if there are no labels in ℱ_1_ at all, then ℱ_2_ must be deleted to match and so we compute the cost of deleting all labels in ℱ_2_. In the second case, label *a* is present in _1_ but not in _2_. Since none of the edit operations insert a label into the label set of a node, all edit sequences must delete *a* and so we add 1 to the distance and remove *a* from consideration moving forward. In the final case, we try both matching the nodes of ℱ_1_ andℱ _2_ containing *a*, as well as deleting *a* from both forests, and we compute and return the minimum cost edit sequence between these two cases. Since *a* is a label of the root of *T*_1_, when trying to match the nodes corresponding to *a*, it is important we delete all nodes above *v*_2_ so that *a* is also a label of *T*_2_. This is why we split ℱ_2_ into *T*_2_(*v*_2_) and *R*(ℱ_2_, *v*_2_). Also one crucial detail of our setup is that by choosing *v*_2_ to be the highest node in *T*_2_ that shares a label with *v*_1_, we can guarantee that during the process of deleting nodes above *v*_2_, we do not accidentally delete a label that we could have kept via a node expansion on *v*_1_. Since *v*_1_ shares no labels with the deleted nodes, we know that it cannot expand to match to any of these soon-to-be-deleted nodes, and we are sure not to delete too many labels in this case.

As mentioned in the previous subsection, by tracing the path which minimizes label deletions through the recursion tree for Equation 1, we may recover the exact minimum cost edit sequence found by `omltd`(*T*_1_, *T*_2_) for two input trees *T*_1_ and *T*_2_, which when applied to the input trees yields the common alignment tree for `omlta` in an additional 𝒪 (*k*) time.

Equation 1 thus provides the starting point and the core structure for our 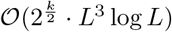-time algorithm for `omlta`. We prove that Equation 1 correctly computes `omltd`(ℱ_1_, ℱ_2_) and that it can be computed in time 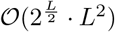, where *L* = |*λ*(ℱ _1_)| + |*λ*(ℱ ^2^)| in Section 4.3. Additional details on how we modify and improve this approach for our final result can be found in Section 4.4. We also provide the proof that computing `omltd` is an NP-hard problem in Section A.3.

### 4.3 Proof of correctness and analysis of asymptotic runtime for the algorithm in Section 4.2

The minimum cost sequence of edits for two forests is typically not unique, since there are often many orderings of the edits that will produce the same transformations. With this in mind, we give a useful observation for computing the `omlta`/`omltd` between two forests in which we have some prior knowledge that a pair of trees, one from each forest, are going to be transformed into the same tree after an optimal sequence of edits. Instead of finding the cost of an optimal edit sequence on the entire forest, we can instead find a minimum-cost sequence of edits on just the trees separately from the rest of the forest and then sum up the computed distances for the pair of trees and the remaining forests. We have implicitly used this idea when building the third case of Equation 1, and now we provide a formal proof of it.

#### Observation 1.

*Given forests* 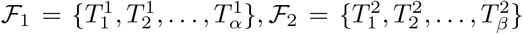 *and a minimum-cost edit sequence S transforming* ℱ_1_ *and* ℱ_2_ *into the same forest, if* 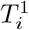 *and* 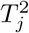 *are transformed into the same forest by S for any i* ∈ [1, *α*], *j* ∈ [1, *β*], *then* `omltd`(ℱ_1_, ℱ_2_) = `omltd` 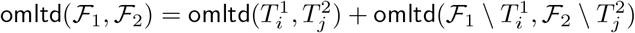.

*Proof*. Consider a minimum cost edit sequence *S* that transforms ℱ_1_ and ℱ_2_ into the same forest such that 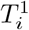 and 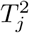 are transformed into the same forest by *S*. Let *c*(*S*) denote the cost, i.e., the number of label deletions, in *S*. Let *S*_*T*_ be the subsequence of *S* that transforms 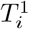 and 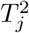 into the same forest, and let *S*_*R*_ be the remaining edits (which transform 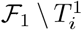 and 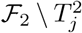 into the same forest). As *S*_*T*_ is a valid edit sequence, `omltd`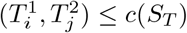. Similarly, `omltd`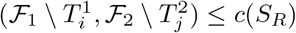. Thus,

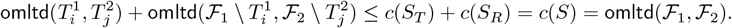

Now for contradiction, assume there is a minimum cost edit sequence 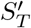 such that 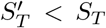 and 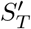 transforms 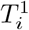 and 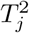 into the same forest. Then 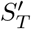, *S*_*R*_ form a valid edit sequence transforming ℱ_1_ and ℱ_2_ into the same forest with 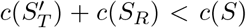, which contradicts *S* being a minimum cost edit sequence. Therefore, no such 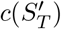 can exist and so `omltd` 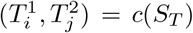 by contradiction. By a similar argument, `omltd` 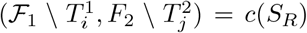, and therefore, `omltd` 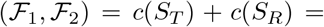 `omltd` 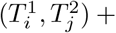 `omltd` 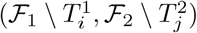.

We make one more useful observation that if two root nodes, one from each forest ℱ _1_ and _2ℱ_, share a label, we can expand the root nodes with this shared label and then compute the rest of the edit distance on the remaining labels in the trees.

#### Observation 2.

*Given trees T*_1_, *T*_2_ *and label a* ∈ *λ*(*r*(*T*_1_)) ∩ *λ*(*r*(*T*_2_)), *then* `omltd`(*T*_1_, *T*_2_) = `omltd`(*T*_1_ \ {*a*}, *T*_2_ \ {*a*})

*Proof*. Consider trees 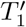 and 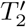 that are the result of performing expansions `expand`(*r*(*T*_1_), {*a*}) and `expand`(*r*(*T*_2_), {*a* }). Any minimum cost edit sequence on *T*_1_ \ {*a*} and *T*_2_ \ {*a*} will be a minimum cost edit sequence on 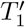 and 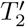 and vise versa. Furthermore, since node expansions are a 0-cost operation, we have that `omltd`(*T*_1_, *T*_2_) = `omltd` 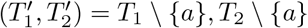.

#### Claim 1.

*Given two multi-label forests* ℱ_1_, ℱ_2_, *Equation 1 correctly computes* `omltd`(ℱ_1_, ℱ_2_).

*Proof*. To compute `omltd`(ℱ_1_, ℱ_2_), recall that by the formalization of Problem 1, we can find a minimum-cost sequence of label deletions, node deletions, and node expansions that transforms _1_ and _2_ into equal forests where the cost of an edit sequence is the number of label deletions performed.

In the first case of Equation 1, when ℱ_1_ is empty, since no allowed edit operations can add any nodes to ℱ_1_, any optimal edit sequence must delete all nodes of ℱ_2_ as well so that both forests are empty after all edits. The number of deleted labels will therefore be |*λ*(ℱ_2_)|, matching the first case of Equation 1.

In the second case, when a mutation label *a* is in ℱ_1_ but not in ℱ_2_, again no allowed edit operations can add *a* to ℱ_2_, so any optimal edit sequence must delete *a*. We may therefore compute `omltd`(ℱ_1_ \ {*a*}, ℱ_2_), the distance between ℱ_1_ without mutation *a* and ℱ_2_, and then add 1 for the deletion of *a*.

In the third case, when mutation label *a* is present in both ℱ_1_ and ℱ_2_, we compute the cost of two possible edit sequences and return the minimum of them. Since all labels only appear once per forest by Definition 1, either both appearances of *a* must be kept in the final versions ofℱ _1_ andℱ _2_ after all edits or both appearances of *a* are deleted. If both are deleted, this adds 2 to the final cost. It is easy to see that any minimum-cost edit sequence transforming ℱ_1_ \ {*a*} and ℱ_2_ \ {*a*} into equal forests will be a minimum-cost edit sequence for ℱ_1_, ℱ_2_ with two extra label deletions to delete the copies of *a*. The final cost of such an edit sequence is `omltd`(ℱ_1_\ {*a*}, *ℱ*_2_ \ {*a*}) + 2, which matches the second parameter of the min function in the third case of Equation 1.

If instead both copies of *a* remain in ℱ_1_, ℱ_2_, note that since *v*_1_ is the root of *T*_1_ ∈ ℱ_1_ and *v*_2_ is the highest node in *T*_2_ sharing a label with *v*_1_, then all labels in an ancestor of *v*_2_ must be deleted in any edit sequence. It may be the case that a minimum-cost edit sequence does not delete the ancestor nodes of *v*_2_ even though they have empty labels, but since node deletions do not contribute to the distance, any such minimum-cost edit sequence may be modified to delete all ancestor nodes of *v*_2_ and the empty-labeled nodes in ℱ_1_ that matched them with no additional cost. Performing these deletions transforms ℱ_2_ into *T*_2_(*v*_2_) ∪ *R*(ℱ_2_, *v*_2_). Furthermore, any optimal edit sequence preserving the *a* labels in *v*_1_ and *v*_2_ must transform *T*_2_(*v*_2_) and *T*_1_ into the same tree. With the above assumption, by Observation 1, we know that

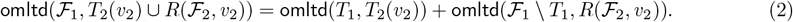

The cost of deleting all ancestors of *v*_2_ is exactly the cost of deleting all labels on the path from *r*(*T*_2_) to *v*_2_ excluding *v*_2_, i.e., |*λ*(*P* (*r*(*T*_2_), *v*_2_))| − |*λ*(*v*_2_)|. Combining this cost with Equality 2 when mutation *a* is preserved by a minimum-cost sequence of edits transforming ℱ_1_ and ℱ_2_ into the same forest yields

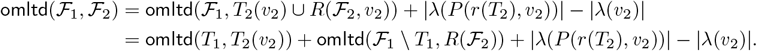

This almost matches the first parameter of the min function in the third case of Equation 1, except that there instead of `omltd`(*T*_1_, *T*_2_(*v*_2_)) we have `omltd`(*T*_1_ \ {*a*}, *T*_2_(*v*_2_) \ {*a*}). However, note that *v*_1_ and *v*_2_ are both root nodes in *T*_1_ and *T*_2_(*v*_2_), respectively, so by Observation 2, `omltd`(*T*_1_, *T*_2_(*v*_2_)) = `omltd`(*T*_1_ \{*a*}, *T*_2_(*v*_2_) \{*a*}). Thus, we correctly compute both the cost of an edit sequence preserving label *a* in both forests and an edit sequence deleting label *a*, and we set the distance to be the minimum of the two in the third case of Equation 1.

#### Claim 2.

*Given two multi-label forests* ℱ_1_, ℱ_2_, `omltd`(ℱ_1_, ℱ_2_) *can be computed in time* 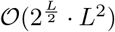, *where L* = |*λ*(ℱ_1_)| + |*λ*(ℱ_2_)|.

*Proof of Claim 2*. By Claim 1, we know an algorithm utilizing Equation 1 will correctly compute `omltd`(ℱ _1_, *ℱ* _2_). We now analyze the runtime complexity of such an algorithm using induction on the number of labels *L* in ℱ_1_ and ℱ_2_. Our base case occurs when ℱ_1_ contains no more labels, which is the first case of Equation 1. In this case, there is no recursion and computing |*λ*(ℱ_2_)| can be done via a simple traversal through ℱ_2_ to count each label, which will take time 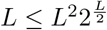.

Now, we define our induction hypothesis to be that `omltd`(ℱ_1_, ℱ_2_) takes time 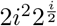 to compute for all forests ℱ_1_, ℱ_2_ with |*λ*(ℱ_1_)|+|*λ*(ℱ_2_)| = *i < L*. Now, consider forests ℱ_1_, ℱ_2_ such that |*λ*(ℱ_1_)|+|*λ*(ℱ_2_)| = *L*. In the second and third cases of Equation 2, we must search through ℱ_2_ to find label *a* if it exists. This can be done again with a simple linear-time search through ℱ_2_ in time *L*. In the second case, *a* is not present in ℱ_2_ and we have a recursive call on `omltd`(ℱ_1_ \ {*a*}, ℱ_2_). Deleting *a* from the label of node *v* can be done in at most *L* time. By the induction hypothesis, the recursive call resolves in time 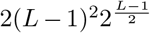, so the total time will be

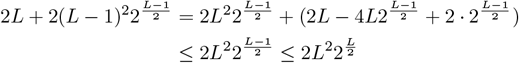

In the third case of Equation 1, we must construct *R*(ℱ_2_, *v*_2_) and remove *a* from ℱ_1_ and ℱ_2_. This can be done in at most 2*L* steps by deleting all labels one at a time in *λ*(*P* (*r, v*_2_)) excluding the labels of *v*_2_ itself, and when a node has its last label deleted, we delete the node itself as well. Finally, there are 3 recursive calls to `omltd`. By the inductive hypothesis, `omltd`(*T*_1_ \ {*a*}, *T*_2_(*v*_2_) \ {*a*}) takes time at most 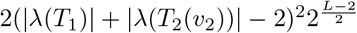 and `omltd`(ℱ \ *T, R*(ℱ_2_, *v*_*2*_)) takes time at most 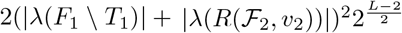 . Note that |*λ*(ℱ_1_ \ *T*_1_)|+|*λ*(*T*_1_)| = |*λ*(ℱ_1_)| and |*λ*(*R*(ℱ_2_, *v*_2_))|+|*λ*(*T*_2_(*v*_2_))| ≤ |*λ*(ℱ_2_)|, so together these recursive calls take time

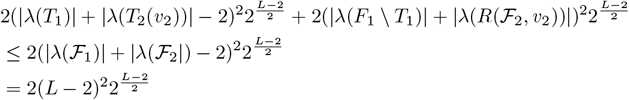

As for the last recursive call `omltd`(ℱ_1_ \ {*a*}, ℱ_2_ \ {*a*}), by the induction hypothesis this takes time 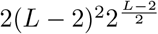 . In total, for the third case we have time

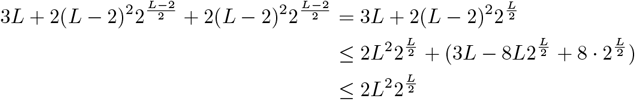

where the last inequality is true for *L* ≥ 2, which must be satisfied in the third case of Equation 1 since both ℱ_1_ and ℱ_2_ contain (a copy of) the label *a*.

Therefore, both recursive cases of Equation 2 are done in time 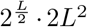 and so, by induction, the entire computation of `omltd`(ℱ_1_, F _2_) can be done in time 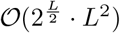.

### 4.4 An asymptotically faster 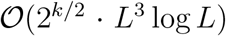-time algorithm for computing omlta/omltd

In this section, we show how to compute `omlta`*/*`omltd` utilizing an improvement to the recursive algorithm defined by Equation 1 such that the runtime depends on *k*, the optimal edit distance between the two input forests. We start by giving the new recursive definition, which takes an additional parameter *k*_*u*_, that can be viewed as an upper bound on *k* that we use to bound the runtime of the recursive computation.

We define the same set of variables as for Equation 1. We let *v*_1_ be the root of any tree *T*_1_ ∈ ℱ_1_, and let *b* ≈ *λ*(*v*_1_). Let *u* be the node in tree *T*_2_ ∈ ℱ_2_ with *b* ∈ *λ*(*u*), if it exists. If such a *u* exists, let *v*_2_ denote the node in the path from *r*(*T*_2_) to *u*, inclusive, that is closest to *r*(*T*_2_) and shares some label with *v*_1_, i.e., *λ*(*v*_1_) ∩ *λ*(*v*_2_) =? ∅. Set *a* to be any mutation in *λ*(*v*_1_) ∩ *λ*(*v*_2_) or just *b* if no matching label for *b* was found in ℱ_2_. We assume implicitly that when the last label of a node is deleted at any step, the now empty-labeled node is deleted as well.

There are two new variables we must define. Let *d*_*T*_ = min{`omltd`(*T*_1_, *T*_2_(*v*_2_)) + |*λ* (*P* (*v*_2_, *r*(*T*_2_))) | − |*λ*(*v*_2_)|, 2} and *d*_ℱ_ = min{`omltd`(ℱ_1_ \{*T*_1_}, ℱ_2_ \{*T*_2_}) +|*λ* (*P* (*v*_2_, *r*(*T*_2_))) | −|*λ*(*v*_2_)|, 2}. *d*_*T*_ and *d*_*F*_ variables correspond to the `omltd` between pairs *T*_1_, *T*_2_(*v*_2_) and ℱ_1_ *T*_1_, *R*(ℱ_2_, *v*_2_), respectively. When these distances are either 0 or 1, the trees and forests are almost exactly matching already and we do not want to make any substantial edits on them. We add two corresponding cases for when either of these distances is small. While computing tree edit distance is NP-hard, checking if two forests are exactly matching (or off by 1 label) can be done in polynomial time dependent on the total number of labels in the forest, and so computing *d*_*T*_ and *d*_ℱ_ does not increase our asymptotic runtime too much.

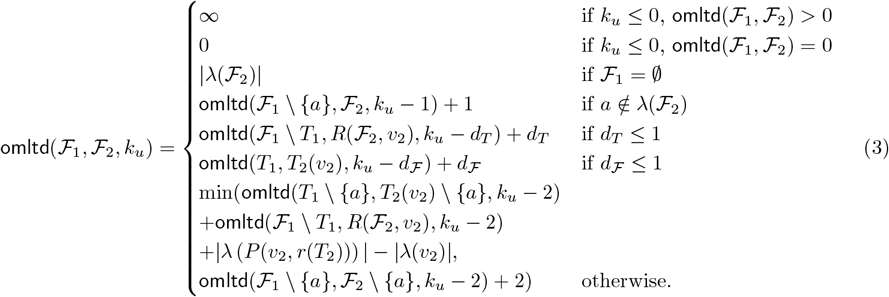

The following two lemmas help prove the correctness of the new fifth and sixth cases in Equation 3. After that, we provide a polynomial-time algorithm for checking if the distance between two forests is at most 1 to show that these cases can be evaluated quickly.

#### Lemma 1.

*Given multi-label forests* ℱ_1_, ℱ_2_ *with trees T*_1_ ≈ ℱ_1_, *T*_2_ ∈ ℱ_2_ *and a* ∈ *λ*(*r*(*T*_1_)) ∩ *λ*(*r*(*T*_2_)). *Let* 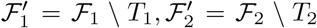. *If d* = `omltd`(*T*_1_, *T*_2_) ≤ 1, *then* `omltd`(ℱ_1_, ℱ_2_) = `omltd` 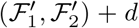 + *d. If d*^*′*^ = `omltd` 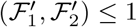, *then* `omltd`(ℱ_1_, ℱ_2_) = `omltd`(*T*_1_, *T*_2_) + *d*^*′*^.

*Proof*. If *d* or *d*^*′*^ are 0, then the lemma statement follows directly from Observation 1.

Consider when *d* = `omltd`(*T*_1_, *T*_2_) = 1. In this case, an optimal edit sequence between *T*_1_ and *T*_2_ must delete a single label *b* present in exactly one of *T*_1_ or *T*_2_. Without loss of generality let *b* ∈ *T*_2_. Now consider a minimum cost edit sequence *S* between ℱ_1_ and ℱ_2_, i.e., a sequence that minimizes the number of label deletions needed to make these two forests equal. If *S* deletes *b*, then *T*_1_ and *T*_2_ already match.

Thus, *S* performs no more label deletions on nodes in *T*_1_ and *T*_2_ and by Observation 1, `omltd` 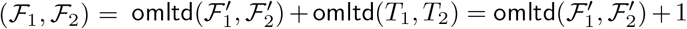. If instead *S* does not delete *b*, then to match the nodes with label *b* in ℱ_1_ and ℱ_2_ together, the roots of *T*_2_ and *T*_1_ can no longer be matched since *b* is in *T*_2_ but not *T*_1_. Since *a* is in the root ofℱ _1_ and ℱ_2_, label *a* must be deleted since these roots are no longer matched together. Since there are two copies of *a, S* must have two label deletions corresponding to deletions of *a*. We may consider a new edit sequence obtaining by modifying *S* to delete both copies of *b* instead and then match *T*_1_ and *T*_2_, which match perfectly after deletion of *b*, and note that the new sequence has the same number of label deletions as *S*. Therefore, any minimum cost edit sequence can be modified to delete *b*, match *T*_1_ and *T*_2_ and so, `omltd` 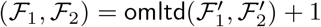.

If `omltd` 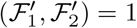, then there must be a label *b* present in exactly one of 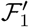 or 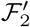 . Without loss of generality let 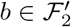 . Consider a minimum cost edit sequence *S* between ℱ_1_ and ℱ_2_. If *S* deletes *b*, then by Observation 1, `omltd`(ℱ_1_, ℱ_2_) = `omltd`(*T*_1_, *T*_2_) + 1. If instead *S* does not delete *b*, then its copy must be in *T*_1_. Similarly to the previous case above, since *a* is in the root of *T*_1_, it must be deleted by *S* in order for the nodes labeled with *b* to match. We consider a new edit sequence obtained by modifying *S* to delete both copies of *b* instead of both copies of *a* and by Observation 2, since *a* labels the root of both *T*_1_ and *T*_2_, the modified edit sequence of *S* still transforms ℱ_1_ and ℱ_2_ into the same forests and requires no additional label deletions. Therefore, any optimal edit sequence can be modified to delete *b* and subsequently match 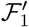 and 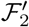 ; therefore, `omltd` 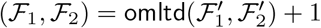.

#### Lemma 2.

*Given multi-label forests* ℱ_1_, ℱ_2_ *with trees T*_1_ ≈ ℱ_1_, *T*_2_ ∈ ℱ_2_ *and a node v* ∈ *T*_2_ *such that*

- *v* ≈ `child`(*r*(*T*_2_)),
- *there exists a label a* ≈ *λ*(*r*(*T*_1_)) ∩ *λ*(*v*),
- |*λ*(*r*(*T*_2_))| = 1.
- *λ*(*r*(*T*_1_)) ∩ *λ*(*r*(*T*_2_)) = ∅

*If* `omltd`(*T*_1_, *T*_2_(*v*)) = 0, *then* `omltd`(ℱ_1_, ℱ_2_) = `omltd`(ℱ_1_ \ *T*_1_, *R*(ℱ_2_, *v*)) + 1. *If* `omltd`(ℱ_1_ \ *T*_1_, *R*(ℱ_2_, *v*)) = 0, *then* `omltd`(ℱ_1_, ℱ_2_) = `omltd`(*T*_1_, *T*_2_(*v*)) + 1.

*Proof*. Since |*λ*(*r*(*T*_2_))| = 1, there is only a single label on *r*(*T*_2_). Let *b* denote this label. Let *S* be a minimum cost edit sequence of ℱ_1_ and ℱ_2_.

If *S* deletes *b*, then it can be modified to delete *r*(*T*_2_) for no cost since empty label deletions are not counted towards `omltd`. If `omltd`(*T*_1_, *T*_2_(*v*)) = 0, then *S* does not have to make any further edits to *T*_1_ and *T*_2_ and so by Observation 1, *mltd*(ℱ_1_, ℱ_2_) = `omltd`(ℱ_1_ \ *T*_1_, *R*(ℱ_2_, *v*)) + `omltd`(*T*_1_, *T*_2_(*v*)) + 1 = `omltd`(ℱ_1_ \ *T*_1_, *R*(ℱ_2_, *v*)) + 1. By a symmetric argument, if `omltd`(ℱ_1_ \ *T*_1_, *R*(ℱ_2_, *v*)) = 0, `omltd`(ℱ_1_, ℱ_2_) = `omltd`(*T*_1_, *T*_2_(*v*)) + 1.

If *S* does not delete *b*, we claim that *S* must delete label *a*. Note that *a* ∈ *λ*(*r*(*T*_1_)) and *b* is the only label of *r*(*T*_2_), which is not included in *λ*(*r*(*T*_1_)) since *λ*(*r*(*T*_1_)) ∩ *λ*(*r*(*T*_2_)) = ∅. Furthermore, *a* is in the label set of *v*, a child of *r*(*T*_2_). The only way for *a* to remain in the forests is if *r*(*T*_2_) is deleted and *v* becomes a root node of its own subtree, but note that deleting *r*(*T*_2_) would require deleting label *b*. By our assumption, *r*(*T*_2_) must not be deleted since *b* is not deleted. Therefore, *a* must be deleted. Since there are 2 label deletions for *a*, we may modify *S* to instead match *a* together and delete both copies of *b* for the same cost. By Observation 2, after deleting *b*, we know that the modified edit sequence of *S* still transforms ℱ_1_, ℱ_2_ into equal forests and no further label deletions need to be added to *S*. Therefore, all edit sequences can be modified to delete *b* instead of *a*. Thus, if `omltd`(*T*_1_, *T*_2_(*v*)) = 0 or `omltd`(ℱ_1_ \ *T*_1_, *R*(ℱ_2_, *v*)) = 0, the lemma statement holds by Observation 1.

#### Lemma 3.

*Given multi-label forests* ℱ_1_, ℱ_2_, min{`omltd`(ℱ_1_, ℱ_2_), 2} *can be determined in* 𝒪 (*L*^2^ log *L*) *time where L* = |*λ*(𝒪_1_)| + |*λ*(𝒪_2_)|.

*Proof*. We now describe an algorithm that computes min{`omltd`(𝒪_1_, 𝒪_2_), 2}.

First, we check if the label sets of 𝒪_1_ and 𝒪_2_ differ by at most 1 label. If they differ by more than 1 label, we may return 2 immediately. If they differ by a single label, we may delete the label. This can be done in 𝒪 (*L*)-time. For the remaining labels present in both forests, if any label *a* in 𝒪_1_ must be deleted, label *a* must also be in 𝒪_2_ since by definition, there is only one copy of each label per multi-label forest. Therefore, if a label shared in both forests is deleted, then `omltd`(𝒪_1_, *F*_2_) ≥ 2. Thus, we can check if 𝒪_1_ and 𝒪_2_ are exactly the same up to node expansions as no labels may be deleted, and we will find max(`omltd`(𝒪_1_, 𝒪_2_), 2). There are many ways to check forest equality, for the sake of completion we outline one below. Define a *branching* node as a node with more than one child and *non-branching* nodes as a node with 1 or 0 children. Note that any node expansion creates a non-branching node, and so when checking if trees are equal up to node expansions, we must check if paths of non-branching nodes can be expanded to become equal. This is done in step 3 of the following routine for checking forest equality.

1. Compute set 𝒫_1_ of paths *P* = (*v*_1_, *v*_2_, …, *v*_*i*_) ≈ ℱ_1_ such that
  - *v*_1_ is the node in *P* closest to the root of the tree containing *P* in ℱ_1_,
  - the parent of *v*_1_ is a branching node or *v*_1_ is the root of the tree containing *P*,
  - *v*_*i*_ is a branching node or a leaf. We call *v*_1_ the *root* of path *P* . We assume that node *v*_1_ has a pointer to its path *P* and vice versa.
2. Sort 𝒫_1_ and 𝒫_2_ lexicographically according to *λ*(*P*) for all *P* ∈ 𝒫_1_ ∪ 𝒫_2_.
3. For each path *P*_1_ ≈ 𝒫_1_:
  a. Use binary search on 𝒫_2_ to find a path *P*_2_ in ℱ_2_ with exactly the same label set as *P*_1_ (i.e., *λ*(*P*_1_) = *λ*(*P*_2_)). If no such path is found, return 2.
  b. Check if *P*_1_ and *P*_2_ are equal up to node expansions. We can do this by checking the labels of the nodes in the paths one at a time starting at the roots of the paths according to Observation 2. If both paths’ roots contain the label, we may remove the label from consideration. Once a root node has all labels removed, we delete it and move to the next node in the path. If the current pair of root nodes considered has no more shared labels and both root node label sets are still non-empty, we return 2.
  c. If *P*_1_ and *P*_2_ are empty after the previous step, we mark these paths as matching by storing a pointer to *P*_2_ in *P*_1_ and move on to the next path of 𝒫_1_.
4. Starting in the root node of a tree *T* in ℱ_1_ we find it’s corresponding path *P* in 𝒫_1_ and the marked matching path *P*^*′*^ in 𝒫_2_. Let *b*_*P*_ be the last node in *P* and let 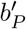 be the last node in *P*^*′*^. Note by Step 1 these are branching nodes or leaves. For every child node *c*_*i*_ of *b*_*P*_ for 1 ≤ *i* ≤ |`child`(*b*_*P*_)| we find it’s corresponding path *P*_*i*_ ∈ 𝒫_1_ and check that the root of the matching path 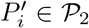 has parent 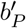 . If the parents of the checked pairs of paths are not *b*_*P*_ and 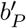, we output 2, otherwise we continue iteratively, performing the same check for the children nodes of the last node in *P*_*i*_ and 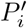 for all 1 ≤ *i* ≤ |`child`(*b*_*P*_)|. We then continue iteratively until we’ve performed these checks for every pair of matching paths. After all paths in tree *T* pass these checks, we move on to a new tree in ℱ_1_ until all paths of ℱ_1_ are checked or 2 is returned.

The first and last step above take 𝒪 (*L*) time and sorting in step 2 can be done in 𝒪 (*L* log *L*) (e.g., via mergesort). The third step dominates the runtime. Performing the binary search on 𝒫_2_ may take time 𝒪 (log *L*). When checking if two paths are equal up to node expansions, each label in ℱ_1_ is only considered during its corresponding path of 𝒫_1_, and it is compared to at most every label of ℱ_2_. Therefore, this step takes time at most 𝒪 (*L*^2^ log *L*). Overall, checking if `omltd`(ℱ_1_, ℱ_2_) ≤ 2 can be done in time 𝒪 (*L*^2^ log *L*).

Finally, we prove the correctness and runtime of an algorithm computing `omltd` according to Equation 3. We begin with the correctness.

#### Lemma 4.

*Given two multi-label forests* ℱ_1_, ℱ_2_, `omltd`(ℱ_1_, ℱ_2_, *k*_*u*_) *for a positive integer k*_*u*_ *computes* min{`omltd`(F_1_, F_2_), *k*_*u*_ + 1}.

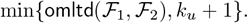

*Proof*. First, we consider when *k*_*u*_ *≥* `omltd`(ℱ_1_, ℱ _2_) and show Equation 3 correctly computes `omltd`(ℱ_1_, *ℱ*_2_) when this inequality holds. As an invariant, we will argue that we maintain that the value passed to parameter *k*_*u*_ is larger than the distance between the two forests of any recursive call we make in any case of Equation 3.

The first three cases of Equation 3 do not have any recursive calls and comprise the base cases.

**Case 1:** In the first case *k*_*u*_ ≤ 0 and `omltd`(ℱ_1_, ℱ_2_) *>* 0. Therefore, the first case can never occur when *k*_*u*_ ≥ `omltd`(ℱ_1_, ℱ_2_).

**Case 2:** The second case is trivially correct. If *k*_*u*_ ≥ `omltd`(ℱ_1_, ℱ_2_) = 0 then 0 is correctly output.

**Case 3:** Similarly, the third case occurs when ℱ_1_ is empty and any optimal edit sequence must delete all of ℱ_2_, which has cost `omltd`(ℱ_1_, ℱ_2_) = `omltd`(∅, ℱ_2_) = |*λ*(ℱ_2_)|, as in Equation 3.

**Case 4:** The correctness of the fourth case follows from Claim 1. However, we must also show that *k*_*u*_ − 1 ≥ `omltd`(ℱ_1_ \ {*a*}, ℱ_2_). By the case constraint, *a* must be deleted by any optimal edit sequence between ℱ_1_ and ℱ_2_ since *a* is only contained in ℱ_1_. Therefore,

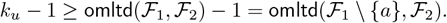

**Case 5:** In the fifth case, we have that *d*_*T*_ ≤ 1 where recall that *d*_*T*_ = min{`omltd`(*T*_1_, *T*_2_(*v*_2_)) + |*λ* (*P* (*v*_2_, *r*(*T*_2_))) | − |*λ*(*v*_2_)|, 2}. Note that since `omltd`(*T*_1_, *T*_2_(*v*_2_)) is non-negative, if *d*_*T*_ ≤ 1, it must be the case that |*λ* (*P* (*v*_2_, *r*(*T*_2_))) | − |*λ*(*v*_2_)| ≤ 1. If |*λ* (*P* (*v*_2_, *r*(*T*_2_))) | − |*λ*(*v*_2_)| = 0, then *v*_2_ is the root of *T*_2_ and so `omltd`(*T*_1_, *T*_2_(*v*_2_)) = `omltd`(*T*_1_, *T*_2_). The correctness of this case then follows from Lemma 1. If instead |*λ* (*P* (*v*_2_, *r*(*T*_2_))) | − |*λ*(*v*_2_)| = 1, then *v*_2_ must be the child of *r*(*T*_2_). Furthermore, since *d*_*T*_ ≤ 1, we must have that `omltd`(*T*_1_, *T*_2_(*v*_2_)) = 0, and so the correctness of this case follows from Lemma 2.

**Case 6:** The correctness of the sixth case follows from an analogous argument to the fifth case.

**Case 7:** The correctness of the final case follows from Claim 1, however, we must make sure that *k*_*u*_ *−* 2 is a correct upper-bound for each recursive call made in this case. Consider the first parameter of the min function. Recall that `omltd`(*T*_1_ \ {*a*}, *T*_2_(*v*_2_) \ {*a*}) + `omltd`(ℱ_1_ \ *T*_1_, *R*(ℱ_2_, *v*_2_)) + |*λ* (*P* (*v*_2_, *r*(*T*_2_))) | − |*λ*(*v*_2_)| computes the cost of a minimum cost edit sequence matching label *a* together in ℱ_1_ and ℱ_2_ by deleting all ancestors of *v*_2_ in ℱ_2_. We know that since we are not in Case 6, *d*_*ℱ*_ = `omltd`(ℱ_1_ \ *T*_1_, *R*(ℱ_2_, *v*_2_)) + |*λ* (*P* (*v*_2_, *r*(*T*_2_))) | − |*λ*(*v*_2_)| ≥ 2. Therefore, if this edit sequence is optimal, we have that

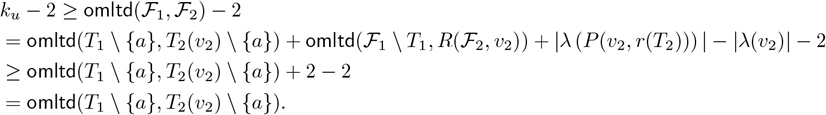

We also know that since we are not in Case 5, *d*_*T*_ = `omltd`(*T*_1_, *T*_2_(*v*_2_)) + |*λ*(*P* (*v*_2_), *r*(*T*_2_))| − |*λ*(*v*_2_)| ≥ 2. An analogous argument shows that *k*_*u*_ *−* 2 ≥ `omltd`(ℱ_1_ *T*_1_, *R*(ℱ _2_, *v*_2_)) as well.

Recall that the second parameter of the min function represents an optimal sequence of edits that deletes label *a* from ℱ_1_ and ℱ_2_. Since *a* is contained in both forests, this requires 2 deletions and so

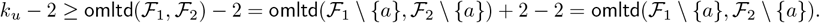

Therefore, we maintain the invariant that *k*_*u*_ ≥ `omltd`(ℱ_1_, ℱ_2_) in any recursive call of Equation 3 if it is true at the first call, and so the algorithm will correctly output `omltd`(ℱ_1_, ℱ_2_) by induction.

If *k*_*u*_ *<* `omltd`(ℱ_1_, ℱ_2_), the correctness of Equation 3 still holds from the previous arguments. However, we may additionally reach Case 1, which returns ∞ or we may compute a number higher than *k*_*u*_. If `omltd`(ℱ_1_, ℱ_2_, *k*_*u*_) computes a value greater than *k*_*u*_, we return *k*_*u*_ + 1, which satisfies the lemma statement.

We now prove the runtime of an algorithm implementing Equation 3 via induction. The proof follows similarly to that of Claim 2.

#### Theorem 1.

*Given two multi-label forests* ℱ_1_, ℱ_2_ *and k* = `omltd`(ℱ_1_, ℱ_2_), *then* `omltd`(ℱ_1_, ℱ_2_) *can be computed in time* 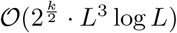.

*Proof*. By Lemma 4, we know that an algorithm utilizing Equation 3 will correctly compute `omltd`(ℱ_1_, ℱ_2_) using *k ≥* `omltd`(ℱ_1_, *ℱ* _2_). We now analyze the runtime complexity of such an algorithm using induction on *k*.

By Lemma 3, we can check the constraints of the fifth and sixth cases of Equation 3 in less than *CL*^2^ log *L* time where *C >* 3 (note the lower bound on 3 is arbitrary and we choose it for convenience later in this proof) is a constant and *L* = |*λ*(ℱ_1_)| + |*λ*(ℱ_2_)|. Now, we define our induction hypothesis to be that `omltd`(ℱ _1_, ℱ _2_, *k*) takes time 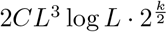.

**Cases 1-3:** Our base case occurs when either of the first three cases of Equation 3 are reached. In any of these cases, there is no recursion. Checking if `omltd`(ℱ_1_, ℱ_2_) = 0 can be done in time *CL*^2^ log *L* by Lemma 3 and computing |*λ*(ℱ_2_)| can be done via a simple traversal through F_ℱ_ to count each label, which will take time 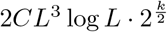in any of these cases.

**Case 4:** In the fourth case, there is a recursive call on `omltd`(ℱ_1_ \ {*a*}, ℱ_2_, *k* − 1), which takes time 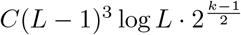 by the induction hypothesis. Finding and deleting label *a* can be done in 2*L* time, and so in total, this step takes

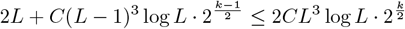

**Case 5:** In this case, we must find label *a* and construct *R*(ℱ_2_, *v*_2_), which can be done in 2*L* time. Additionally, we must compute *d*_*T*_ . By Lemma 3, this can be done in time at most *C*(|*λ*(*T*_1_)|+|*λ*(*T*_2_)|)^2^ log *L*. By the induction hypothesis, the recursive call takes time at most 2*C*(|*λ*(ℱ_1_ \{*T*_1_ }|+|*λ*(ℱ_2_)\{*T*_2_ }|)^3^ log 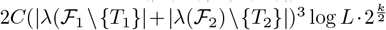·

Let *S* = |*λ*(*T*_1_)| + |*λ*(*T*_2_)|. The total time of this step is then at most

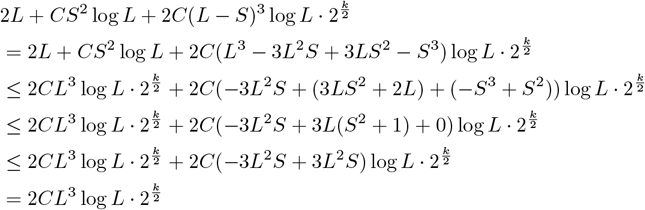

where note the second last line follows since *S*^2^ + 1 ≤ *LS* since *T*_1_ and *T*_2_ contain at least label *a* each so *S >* 0 and *L > S* (if *L* = *S*, we can skip the recursive call as the remaining forests would be empty).

**Case 6:** This case follows from a symmetric argument to case 5.

**Case 7:** This is the final case. As for Equation 1, we must construct *R*(ℱ_2_, *v*_2_) and remove *a* from ℱ_1_ and ℱ_2_. This can be done in at most 2*L* steps by deleting all labels one at a time in *λ*(*P* (*r, v*_2_)) excluding the labels of *v*_2_ itself, and when a node has its last label deleted, we delete the node itself as well. We additionally must compute *d*_*T*_ and *d*_*ℱ*_ to check if we indeed fall into any of the previous cases, which takes time at most *CL*^2^ log *L*. There are three recursive calls we consider. Let *S* = |*λ*(*T*_1_)| + |*λ*(*T*_2_)|. By the inductive hypothesis, `omltd`(*T* \ {*a*}, *T* (*v*) \ {*a*}, *k* − 2) takes time at most 2*C*(*S* − 2)^3^ log 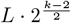 and `omltd`(ℱ_1_ \ *T*_1_, *R*(ℱ_2_, *v*_2_), *k* − 2) takes time at most 2*C*(*L* − *S*)^3^ log 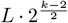 . Together, these recursive calls take time

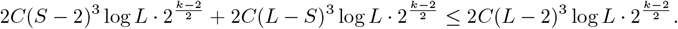

As for the last recursive call, `omltd`(ℱ_1_ \ {*a*}, ℱ_2_ \ {*a*}, *k* − 2) takes time at most 2*C*(*L* − 2) log 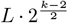 by the inductive hypothesis. We therefore have total time at most,

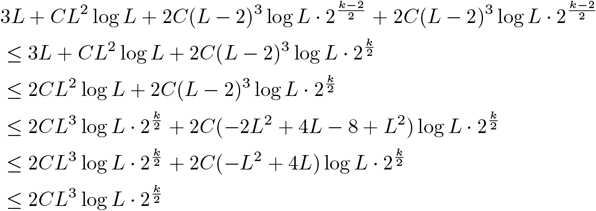

where the last inequality follows since *L*^2^ ≥ 4*L* for all *L* ≥ 4. We know that *L* is at least 4 since otherwise we would have fallen into case 5 or case 6 as there would be no labels in ℱ_1_ andℱ _2_ besides both copies of *a* and potentially one other label.

Therefore, all cases take time at most 2*CL*^3^ log 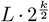 and so by induction, `omltd`(ℱ_1_, ℱ_2_, *k*) takes time 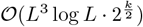.

Given an upper bound *B* ≥ `omltd`(ℱ_1_, ℱ_2_), Theorem 1 provides a very fast bound 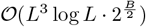 for `omlta`*/*`omltd` if `omltd`(ℱ _1_, *ℱ* _2_) ≤*B*, otherwise it reports that there is no valid solution. However, for real data sets we cannot know in advance what value to assign to *B*. We run our algorithm multiple times incrementing *B* by 1 at each iteration and starting at *B* = 1 until our algorithm outputs a valid solution. Let *k* = `omltd`(ℱ_1_, ℱ_2_). The total runtime to compute `omltd`(ℱ_1_, ℱ_2_) considering these *k* iterations will be:

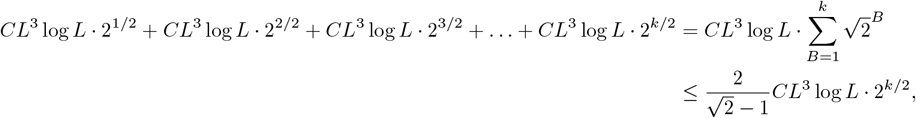

where the above inequality follows from the geometric series formula.

#### Obtaining omlta from omltd calculation

As shown, Equation 3 provides a recursive computation of `omltd` that can be used to find the minimum number of label deletions for two input trees *T*_1_ and *T*_2_. In other words, in the recursion tree for this computation, there exists at least one optimal branch that can be used to compute `omltd`. If we keep track of the optimal branch of the recursive algorithm in the recursion tree, we may additionally construct a minimum cost edit sequence that constructs the `omlta` alignment tree. We describe how to do so informally in the following paragraph using the notation of Equation 3.

While following the optimal branch, we add label deletions to our minimum cost edit sequence corresponding to removed labels in cases 3, 4, 5, 6, where the latter two cases only have a label deletion if *d*_*T*_ = 1 or *d* _*ℱ*_= 1. In case 7, there are two subcases. Either we delete label *a* in both subtrees, which if optimal, we add as a label deletion for *T*_1_ and *T*_2_ in our minimum cost edit sequence, or we delete all nodes and labels in the path leading to node *v*_2_, which again we add to our edit sequence if optimal. In the latter subcase, we choose to retain label *a* in *T*_1_ and *T*_2_, which is maintained as we continue down the optimal branch, so we may add a node expansion for *v*_1_ and *v*_2_ to our minimum cost edit sequence to separate label *a* from *v*_1_ and *v*_2_ into a pair of matching nodes in the final alignment tree. Finally, after reaching the bottom of the optimal recursion tree branch, we obtain all label deletions and node expansions. We may then finish our minimum cost edit sequence by adding any node deletions needed for any nodes whose entire label set is deleted in the minimum cost edit sequence. By performing the resulting edit sequence on *T*_1_ and *T*_2_, both are transformed into the alignment tree for `omlta`.

## Acknowledgments

This research was supported in part by the Intramural Research Program of the National Institutes of Health (NIH). The contributions of the NIH authors were made as part of their official duties as NIH federal employees, are in compliance with agency policy requirements, and are considered Works of the United States Government. However, the findings and conclusions presented in this paper are those of the authors and do not necessarily reflect the views of the NIH or the U.S. Department of Health and Human Services. This work utilized the computational resources of the NIH HPC Biowulf cluster (https://hpc.nih.gov).

## Supplementary Materials

## A Supplementary Materials

### A.1 Related works on unordered tree edit distance

The first algorithm of (5) is the most similar to our own algorithm both in terms of runtime (many of the related works we list below depend on other parameters rather than *k*) and technical approach. To find the maximum-size tree alignment, the algorithm of (5) constructs a minimum-edit mapping between the nodes of each input tree, allowing nodes to map to and from an empty set, which represents the edits of insertions and deletions, while nodes with different label sets mapped together represent label substitution edits. Such a mapping immediately provides the maximum tree alignment of the input trees and can easily be used to find the tree edit distance as well.

When considering a pair of mapped branching nodes (nodes with degree greater than 1), the algorithm of (5) reduces the problem of finding a mapping between the children of these nodes to the well-studied *maximum weighted bipartite matching* (66) problem by considering the set of children of each node in the pair to be the two parts in a bipartite graph and then setting edge weights between two nodes in this graph according to the number of edits in a minimum-edit mapping between the subtrees of these nodes in the original input trees. Observe that this is an inherently recursive approach, the reduction to maximum weighted bipartite matching requires solving a smaller instance of the mapping problem. Our algorithm utilizes a similar top-down recursive approach, but instead of mapping nodes to nodes, we focus on mapping label to label. In this way, we are able to simplify branching node mappings by relying on the uniqueness of labels: we can either map a label of one input tree to its singular copy in the other tree, or we have to delete that label. By carefully arguing that we only have to make such a choice *k* times, we are able to avoid relying on the reduction to bipartite matching while simultaneously improving the runtime, reducing the exponential factor from 2.62^*k*^ to 2^*k/*2^. Furthermore, we are easily able to incorporate the node expansion edit, an edit that as argued above, is useful for correctly capturing dissimilarity in clonal trees.

Other parameterized tree edit distance algorithms have been considered as well; Akutsu et al., 2012 (32) provide an 𝒪 (1.26^*m*^)-time algorithm where *m* is the total number of nodes across both input trees and an 𝒪 ((1 + *ϵ*)^*m*^)-time algorithm for trees of bounded degree for any *ϵ >* 0. Instead of parameterizing on the total number of nodes, Akutsu et al., 2013 (33) consider the number of branching nodes, nodes in either tree with more than one child, and give an 𝒪 (2^*b*^*nd*)-time algorithm where *b* is the number of branching nodes and *d* is the maximum degree of either input tree. Two works (34; 35), explore an approach based on a reduction from tree edit distance to another well-known NP-hard problem, the maximum vertex weighted clique problem. While still working in exponential time, both papers provide practical results for small enough tree instances.

### A.2 Maximum agreement subtree is a different problem from omlta

In the Maximum Agreement Subtree problem (36; 37; 38), a set of trees *T*_1_, *T*_2_, …, *T*_*m*_ are given as input such that each tree *T*_*i*_ has a set of *n* leaf nodes labeled from a label set *L* of size *n* such that the leaf labels of *T*_*i*_ partition *L* for each 1 ≤ *i ≤ m* (internal nodes are not labeled), and the goal is to find the largest label subset *L*^*′*^ ⊆ *L* such that the minimal spanning tree of *T*_*i*_ and *T*_*j*_ for all 1 ≤ *i, j ≤ m* containing label set *L*^*′*^ is equal. Maximal agreement subtree has been studied and has applications in bioinformatics for comparing phylogenetic trees. If `omlta` could be efficiently reduced to maximal agreement subtree, then any useful results would directly transfer to our new problem.

One possible idea towards a reduction from `omlta` would be to take each input tree *T*, which may have internal node labels, perform node expansions (which have no cost in `omlta`) until each node of *T* only has a single label, and then append a new leaf node to each node of the original tree and shift labels to the appended leaf nodes. The resulting set of trees would be of the form required for maximum agreement subtree. The main issue with this reduction is choosing how to expand nodes with multiple labels. Recall that when node expansions occur, we replace a node with a pair of nodes, a parent and child, whose label sets partition the original node’s label set. Since we work with rooted trees, this means that each node expansion requires choosing an ordering for the label set (which labels will move higher in the tree and which will move lower in the tree). This ordering can change the size of the label set in the solution of the maximum agreement subtree instance we obtain. It is therefore impossible to consistently utilize such a reduction without specifying some ordering technique. However, choosing an ordering that provides consistent structure in the maximum agreement subtree problem is non-trivial. If we aim to choose the expansion ordering that maximizes the label set in the solution of the maximum agreement subtree instance, this may require finding one ordering out of an exponential number of orderings. It is unlikely that a polynomial-time algorithm for finding such an ordering can be found.

Interestingly, even for the trees with a single label per node, the above reduction can result in a maximum agreement subtree whose size is substantially different from the `omlta` tree. Consider, for example, trees *T*_1_ with a root node labeled with *a* and its single child labeled with *b*, and *T*_2_ with an unlabeled root and its two children respectively labeled with *a* and *b*. The `omltd` between *T*_1_ and *T*_2_ is 1, which is the maximum distance possible for instances with only two labels. When this reduction approach to maximum agreement subtree is applied to *T*_1_, *T*_1_ is transformed into a tree isomorphic to *T*_2_ - which is already in the desired format - thus their maximum agreement subtree is *T*_2_ itself, with 0 (minimum possible) label deletions. This example can be extended to larger trees in which no leaf has a sibling and the leaf and its parent each has a single label. The `omltd` between such a tree and the tree obtained by applying the above reduction to it would be equal to half of the total number of its labels (other measures for comparing tumor phylogenies, including ancestor-descendant or different-lineage accuracy would also be high), while no labels would need to be deleted for the corresponding maximum agreement subtree instance.

For these reasons and more (which we do not expand on for the sake of brevity), `omlta` is a very different problem from maximum agreement subtree. As a result, while the maximum agreement subtree is a suitable approach for species phylogenies, it is not designed for, or used on tumor phylogenies. The most efficient algorithms for computing the maximum agreement subtree have a running time exponential with the maximum degree of the input trees (36). Even if the above reduction could be completed correctly, it would produce trees with node degrees ≥ the number of labels which could yield high runtimes with maximum agreement subtree algorithms.

Given trees *T*_1_, *T*_2_ and label *a* ∈ *λ*(*r*(*T*_1_)) ∩ *λ*(*r*(*T*_2_)), then `omltd`(*T*_1_, *T*_2_) = `omltd`(*T*_1_ \ {*a*}, *T*_2_ \ {*a*}) By Claim 1, we know an algorithm utilizing Equation 1 will correctly compute `omltd`(ℱ_1_, ℱ_2_). We now analyze the runtime complexity of such an algorithm using induction on the number of labels *L* in ℱ_1_ and ℱ_2_. Our base case occurs when ℱ_1_ contains no more labels, which is the first case of Equation 1. In this case, there is no recursion and computing |*λ*(ℱ_2_)| can be done via a simple traversal through ℱ_2_ to count each label, which will take time 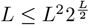.

### A.3 Computational Hardness of omlta/omltd

We show that the `omlta` and `omltd` problems are NP-hard by reducing the **Exact Cover by 3-Sets** (X3C) problem to `omltd`. Exact Cover by 3-Set is well-known to be NP-Complete (67).

#### Definition 3

(Exact Cover by 3-Sets). *Given a finite set* 𝒰 *with* | 𝒰| = *n* = 3*q and a collection* 𝒮 *of 3-element subsets of* 𝒮, *the* Exact Cover by 3-Sets *(X3C) problem is to determine if there exists an* exact cover, *a sub-collection* 𝒮*′ of* 𝒮 *such that every element of* 𝒰 *is in exactly one set of* 𝒮*′*.

Let 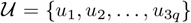 and without loss of generality, let 𝒮 = { 𝒮_1_, 𝒮_2_, …, 𝒮_*m*_} with *m > q*. Define tree 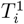 for 1 ≤ *i* ≤ *m* to be a path of length ℓ = 2*q* connected to three paths of length ℓ. The label set of the *j*th node for 1 ≤ *j* ≤ ℓ on the first path are *α*_*i,j*_ = {*a*_*i*,1,*j*_, *a*_*i*,2,*j*_, …, *a*_*i,m,j*_} where each *a*_*i,k,j*_ is a distinct character not in 𝒰 such that *a*_*i,k,j*_ = *a*_*i*_*′*_,*k*_*′*_,*j*_*′* if and only if *i* = *i*^*′*^, *j* = *j*^*′*^, *k* = *k*^*′*^. We call this path the “*α*-path” of *T*_*i*_. Let 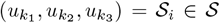 be the *i*th set in 𝒮 ; this is an unordered set, but the order of the numbers 1,2,3 in the subscripts matters for what follows. The label set of the *j*th node on the first of the three remaining paths for 1 ≤ *j* ≤ ℓ is a singleton 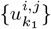. The other two paths are defined analogously for *k*_2_ and *k*_3_. We refer to these paths as the “*u*-paths” of 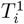. Similarly to the *α*-paths, each label 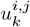 is distinct within a tree and not contained in 𝒰. Let 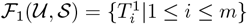.

In the construction above, we have intentionally createdℱ _1_ so that each set 𝒮_*i*_ ∈ *𝒮* has a corresponding tree 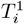 and within each tree 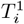, there are exactly three u-paths in one-to-one correspondence with the three elements of 𝒮_*i*_. The lengths of the u-paths are each ℓ = 2*q*, which is exactly 2/3 the number of elements of𝒰 . By making the construction in this way, when we later align ℱ_1_ with a second forest described in the following paragraph, we are able to use the alignment to reason about the original X3C instance in our reduction.

Define tree 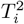 for 1 ≤ *i* ≤ *q* to be a single node connected to three paths of length ℓ. Let the label of the root node of 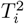 be *β*_*i*,1_ where *β*_*i,j*_ = {*a*_1,*i,j*_, *a*_2,*i,j*_, …, *a*_*m,i,j*_}. The label set of the *j*th node on the first of three paths for 1 ≤ *j* ≤ ℓ is 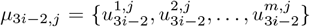 . The other two paths are defined analogously with 3*i*− 1 and 3*i*, respectively. We refer to these paths as the “*u*-paths” of 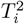 . Define another tree 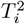 for *q < i* ≤ *m* to be a path of length ℓ. The label set of the *j*th node for 1 ≤ *j* ≤ ℓ on this path is *β*_*i,j*_. We call these paths “*β*-paths.” Let 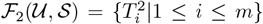. Observe that each of the 3*q* elements of 𝒰 has a corresponding *u*-path in ℱ_2_, distributed across *q* trees.

See Supplemental Figure S1 for an example of these forests for instance (𝒰, 𝒮).

Conceptually, we have built these trees such that if a set *S*_*i*_ is in the solution of an Exact Cover by 3-Set instance, then when solving `omlta` on *F*_1_ and *F*_2_, tree 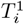 may retain a substantial number of its labels by matching them to those in *F*_2_. Similarly, if there is no solution to the instance, then a large amount of labels of 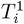 must be deleted. We show formally that we are able to distinguish whether or not such a solution exists by computing the `omltd` for the constructed forests *F*_1_ and *F*_2_ for the X3C instance and determining if this distance is larger than a specific threshold or lower than a specific threshold, given below.

**Figure S1:**
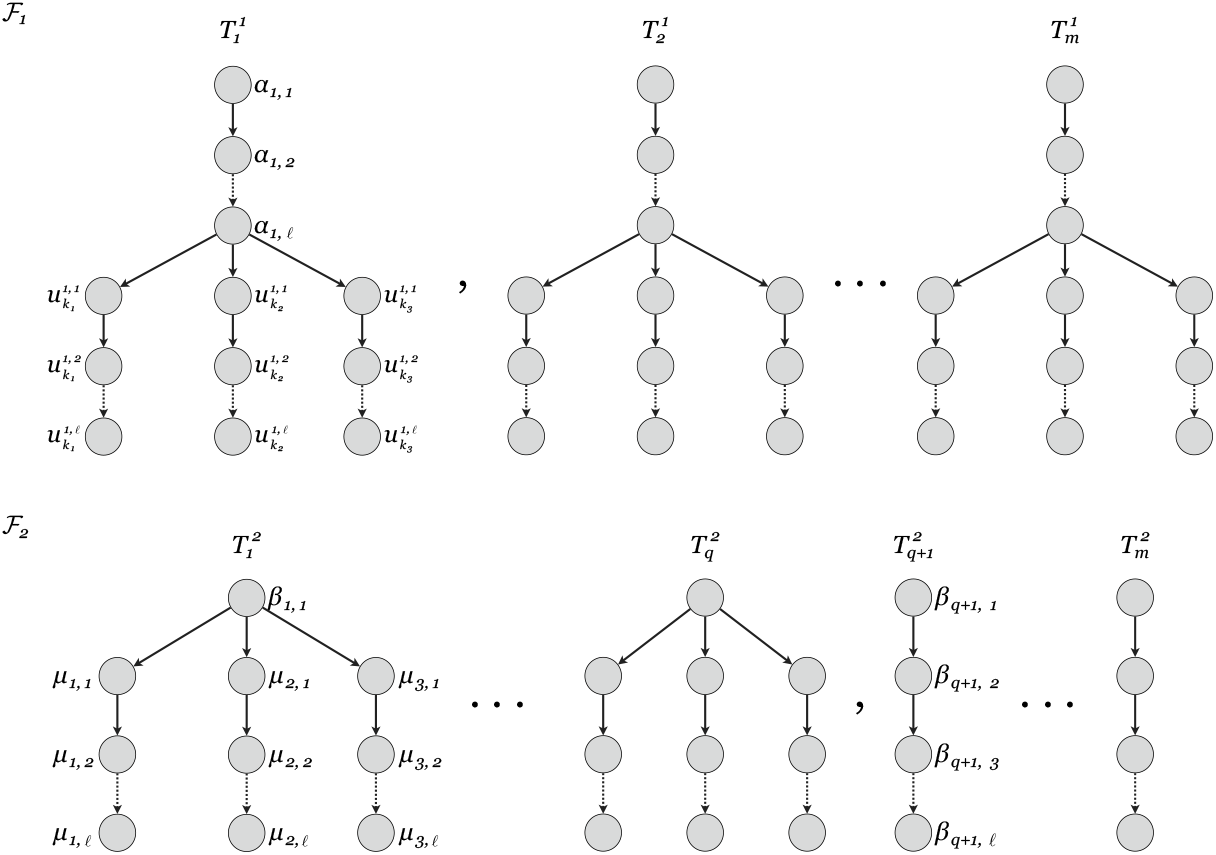
Example of forests ℱ_1_(𝒰, *𝒮*) and ℱ_2_(𝒰, *𝒮*) for X3C instance (𝒰, *𝒮*). In the *u*-paths, what may look like the bottom-most edges are actually dotted lines (like an ellipsis) symbolizing many edges, not solid lines representing single edges. We do not show the excluded edges for the sake of compactness, but they follow exactly according to our construction of ℱ_1_ and ℱ_2_.

#### Lemma 5.

*Given an X3C instance* (𝒰, 𝒮) *with* |𝒮| = *m*, |𝒰| = 3*q*, ℓ = 2*q, and m > q, if there is an exact cover for* (𝒰, 𝒮), *then* `omltd`(ℱ_1_(𝒰, 𝒮), ℱ_2_(𝒰, 𝒮)) ≤ *q*(ℓ + 1)*m* + 3*q*(*m* − 1)ℓ + (*m* − *q*)(2*m* − 2)ℓ + 3(*m* − *q*)ℓ.

*Proof*. Let ℱ_1_ denote *ℱ*_1_(𝒰, 𝒮) and let ℱ_2_ denote ℱ_2_(𝒰, 𝒮). First, if there is an exact cover, there is some sub-collection 𝒮*′* such that every element *u* ∈ 𝒰 occurs exactly once across all sets of 𝒮*′*. In other words, there is a bijection between elements in sets of 𝒮*′* and 𝒰. Therefore, there is a subforest 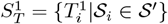 of trees in ℱ_1_ such that there is a bijection between the *u*-paths of 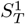 and the *u*-paths of 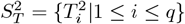. Using this fact, we now describe a sequence of deletions that transform ℱ_1_(𝒰, 𝒮) and ℱ_2_(𝒰, *𝒮*) into the same forest where the number of deletions needed matches the lemma statement.

Note that the *j*th node *v*_1_ on a *u*-path of tree 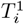 has a single label 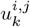, and this label is shared by the *j*th node of a *u*-path in tree 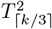 . These *u*-paths may be determined by the bijection between elements of 𝒮*′* and 𝒰. Therefore, we may delete the *α*-paths of all trees in 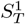, the root nodes of all trees in 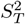, and *m* − 1 labels per remaining node in the *u*-paths of 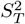 to transform the trees of 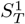 and 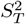 into equal forests. Since |*α*_*i,j*_| = *m*, |*β*_*i,j*_| = *m*, and the length of paths is ℓ, this requires *qm*ℓ, *qm* and 3*q*(*m* 1)ℓ label deletions, respectively.

There are *m* − *q* remaining trees in ℱ_1_ and ℱ_2_ that have been unchanged. The *j*th node in an *α*-path of 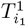 and the *j*th node in a *β*-path of tree 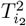 for *q < i*_2_ ≤ *m* share exactly one label 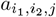. By deleting all the *u*-paths of the remaining *m* − *q* trees in ℱ_1_, we can form a bijection of the remaining *α*-paths of ℱ_1_ and the *β*-paths ofℱ _2_. Deleting all labels on *u*-paths requires 3(*m ™q*)ℓ label deletions and deleting all non-matched labels of the *α* and *β*-paths requires (*m™ q*)(2(*m ™*1))ℓ label deletions since each node has *m* labels and all but 1 of the labels is deleted. The resulting forests are now equal. Since the above edit sequence transforms F_1_ and F_2_ into the same forest, we have that

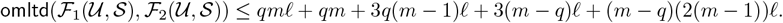

Reordering the terms yields the desired bound.

#### Lemma 6.

*Given an X3C instance* (𝒰, 𝒮) *with* | 𝒮 | = *m*, | 𝒰 | = 3*q*, ℓ = 2*q, and m > q if* `omltd`(ℱ_1_(𝒰, 𝒮), ℱ_2_(𝒰, 𝒮)) ≤ *q*(ℓ + 1)*m* + 3*q*(*m* − 1)ℓ +(*m* − *q*)(2*m* − 2)ℓ + 3(*m* − *q*)ℓ, *then there is an exact cover for* (𝒰, 𝒮).

*Proof*. Let *f* = *q*(ℓ + 1)*m* + 3*q*(*m* − 1)ℓ + (*m* − *q*)(2*m* − 2)ℓ + 3(*m* − *q*)ℓ. Let ℱ_1_ denote *F*_1_(𝒰, 𝒮) and let ℱ_2_ denote ℱ_2_(𝒰, 𝒮). Let ℰ be an edit sequence with at most *f* label deletions that transforms ℱ_1_ and ℱ_2_ into the same forests. We wish to show that if such an ℰ exists, then there must be a bijection between the *u*-paths of ℱ_2_ with the *u*-paths of exactly *q* trees of ℱ_1_ corresponding to a bijection between elements of *u* and a sub-collection of 𝒮, respectively. Thus, if we show such a bijection, then there is an exact cover for (𝒰, 𝒮).

First, we discuss all the possible matching labels between nodes of ℱ_1_ and ℱ_2_. Observe that the *j*th node of an *α*-path in a tree 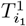 for 1 ≤ *i*_1_ ≤ *m* and the *j*th node of a *β*-path in 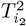 with *q < i*_2_ ≤ share exactly label 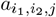. Similarly, the first node of an *α*-path in 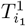 for 1 ≤ *i*_1_ ≤ *m* and the root of 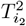 for 1 ≤ *i*_2_ ≤ *q* share exactly label 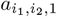. Any nodes of an *α*-path, *β*-path, or root of a tree 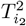 with 1 ≤ *i*_2_ *< q* do not share any labels with other nodes beyond the above cases.

Furthermore, the *j*th node of a *u*-path in 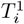 has only a single label 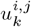 and the *j*th node of a *u*-path 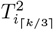 has label 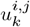 as well. These are the only shared labels between nodes of *u*-paths in ℱ_1_ and ℱ_2_.

Let *d* be the number of *β*-paths who have the labels on all their nodes deleted by ℰ. As previously discussed, *α*-paths only may be matched with *β*-paths of 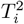 for *q < i* ≤ *m* and root nodes of 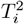 for 1 ≤ *i* ≤ *q*. Since there are exactly *m α*-paths in ℱ_1_ (1 per tree), then there must also be *d α*-paths whose labels are completely deleted in ℱ_1_. Since *α* and *β*-paths have length ℓ and each node has *m* labels, this corresponds to 2*dm*ℓ deletions.

For each remaining (*m*− *q* −*d*) *β*-path, the nodes in the path must be matched to an *α*-path. ℰ must delete all but one label per node on these *β* and *α*-paths since nodes in *α* and *β*-paths only share a single label as discussed previously. Therefore, the remaining *β*-paths and the *α*-paths whose labels are matched correspond to another set of exactly (*m ™q ™ d*)(2*m ™* 2)ℓ deletions.

So far, (*m™ q*) of the *α*-paths have had their labels matched or deleted. For the remaining *q* of them, all but the first of the nodes in the *α*-path may be matched to the root of a tree 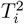 for 1≤ *i < q*. As discussed above, only a single label is shared between these nodes and so, this corresponds to a total of *q*(*m*ℓ + *m* − 2) label deletions.

Next we consider the 3*q u*-paths in ℱ_2_. Since each of the ℓ nodes in a *u*-path of ℱ_2_ has *m* labels and shares a single label with a node in a *u*-path of ℱ_1_ and shares no labels with any nodes outside of a *u*-path, there must be 3*q*(*m* − 1)ℓ label deletions to match the *u*-paths of ℱ_2_ with ℱ_1_. All the labels in the remaining *u*-paths of ℱ_1_ with no matching path in ℱ_2_ must be deleted, which corresponds to 3(*m q*)ℓ label deletions.

In total, ℰ must make a number of label deletions more than

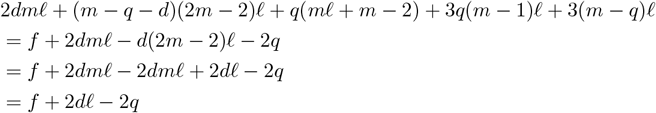

Recall that ℓ *> q* and so 2*d*ℓ *>* 2*q* if *d >* 0. Thus, *d* must be 0 since ℰ makes at most *f* label deletions, and so, no *β*-paths in ℱ_2_ have all their labels deleted by ℰ.

Now, let *d*^*′*^ be the number of *u*-paths in ℱ_2_ whose labels on all their nodes are deleted by ℰ. Since each node in *u*-path of ℱ_2_ has *m* labels and such a path has length ℓ, this corresponds to *d*^*′*^*m*ℓ deletions. Since each of the ℓ nodes in a *u*-path of ℱ_2_ has *m* labels and shares a single label with a node in a *u*-path of ℱ_1_ and shares no labels with any nodes outside of a *u*-path, there must be (3*q* − *d*^*′*^)(*m* − 1)ℓ label deletions to match the remaining *u*-paths of ℱ_2_ with ℱ_1_. All the labels in the remaining *u*-paths of ℱ_1_ with no matching path in F_2_ must be deleted, which corresponds to 3(*m* − *q*)ℓ + *d*^*′*^ℓ label deletions.

As before, to match all *β*-paths in ℱ_1_ with their corresponding *α*-paths in ℱ_2_ requires (*m* − *q*)(2*m* − 2)ℓ label deletions. Similarly, to match all remaining *α*-paths of ℱ_2_ with a root node of 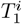 for 1 *< i < q* requires *q*(ℓ + 1)*m* − 2*q* label deletions. The total number of labels deleted by ℰ is therefore

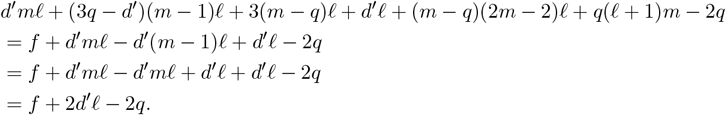

Again, since ℓ *> q*, we have that 2*d*^*′*^ℓ *>* 2*q* if *d*^*′*^ *>* 0. Thus, *d*^*′*^ must be 0 since ℰ makes at most *f* label deletions, and so, no *u*-paths of ℱ_2_ have their labels fully deleted by ℰ.

Since a *β*-path of ℱ_2_ can only be matched to a *α*-path of ℱ_2_ and we’ve shown that no *β*-paths have all their labels deleted, then *m* − *q* trees of ℱ_2_ must have their *u*-paths deleted to match their *α*-paths to the *β*-paths of ℱ_2_. The remaining *q* trees have exactly 3*q u*-paths, which must be matched to the 3*q u*-paths of ℱ_2_ since we’ve also shown that no *u*-path of ℱ_2_ has all its labels deleted. Therefore, there is a bijection between the *u* paths of a collection of *q* trees of Fℱ_1_ and the *u*-paths of ℱ_2_. Thus, there is an exact cover for (𝒰, 𝒰).

#### Theorem 2.

`omlta`*/*`omltd` *computation is NP-hard*.

*Proof*. By Lemma 5 and Lemma 6, the above tree construction is a reduction from NP-Complete problem Exact Cover by 3-Sets to `omltd`.

### A.4 Additional figures on trees inferred by PairTree, CONIPHER and their omlta from NSCLC cohort

**Figure S2:**
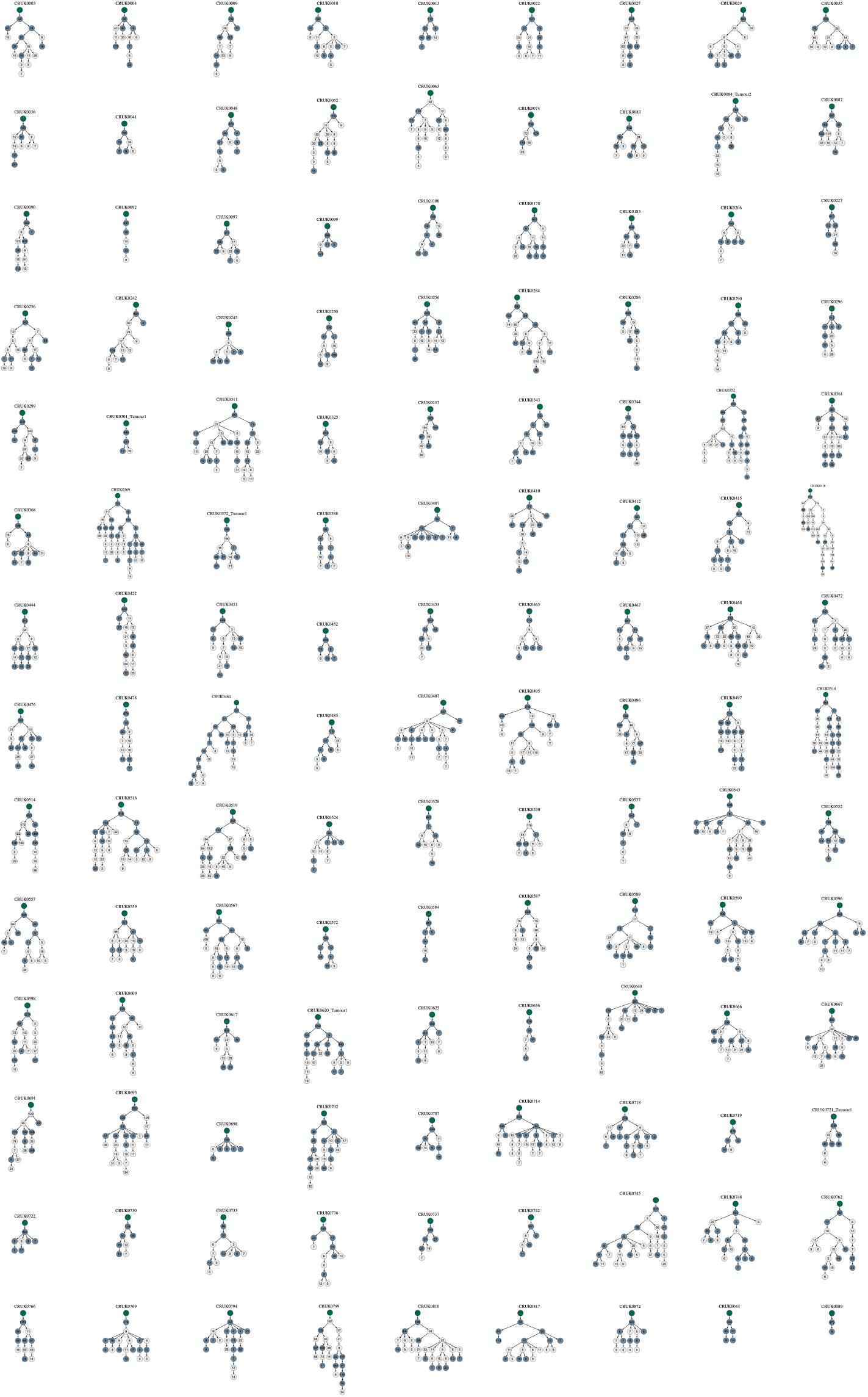
`CONIPHER` trees for each case of the NSCLC cohort. Nodes are labeled by the number of mutations in the cluster and colored green if that node/mutational cluster is preserved in the alignment with the corresponding `PairTree` tree.

**Figure S3:**
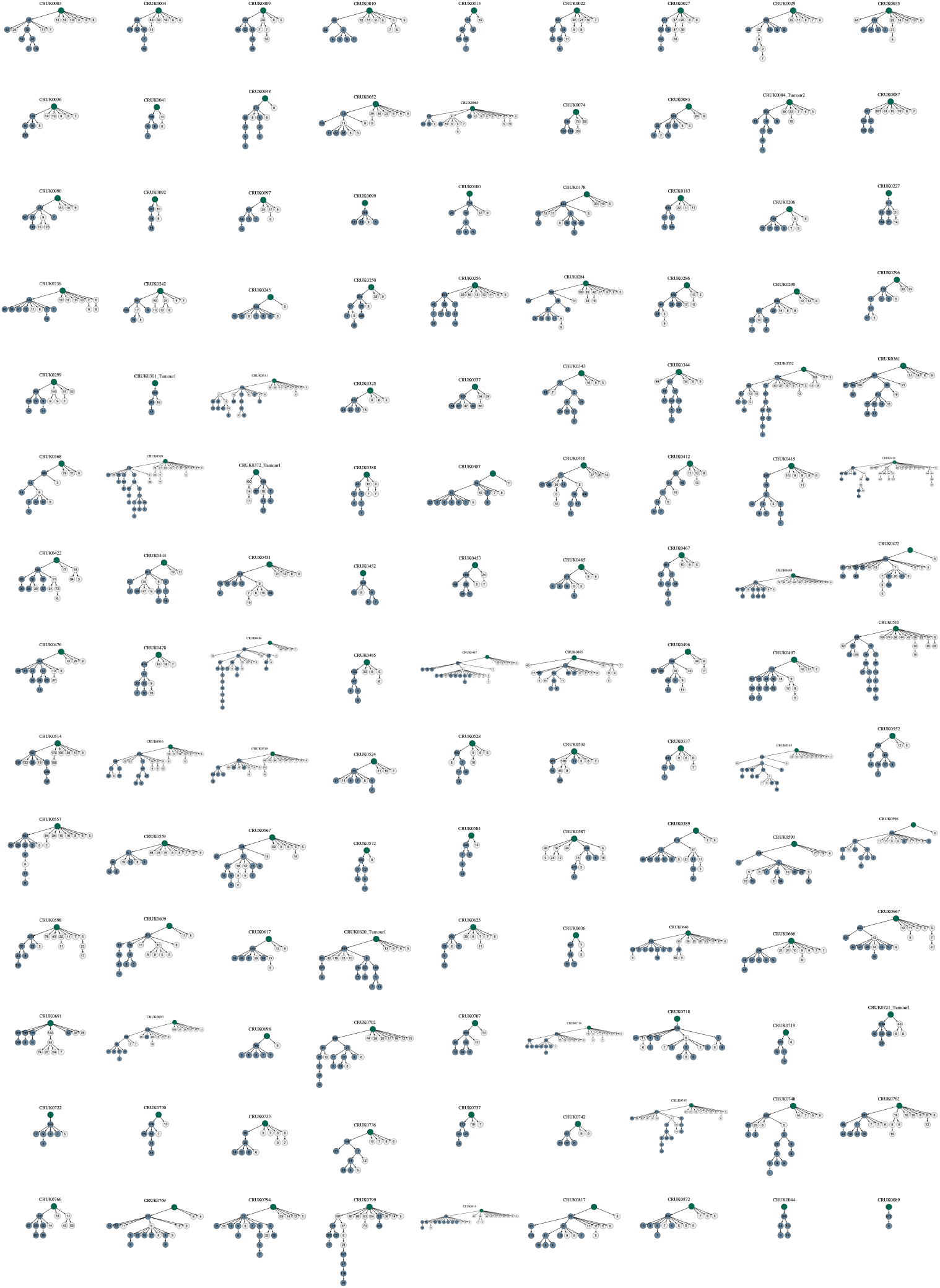
`PairTree` trees `CONIPHER` trees for each case of the NSCLC cohort. Nodes are labeled by the number of mutations in the cluster and colored green if that node/mutational cluster is preserved in the alignment with the corresponding `CONIPHER` tree.

**Figure S4:**
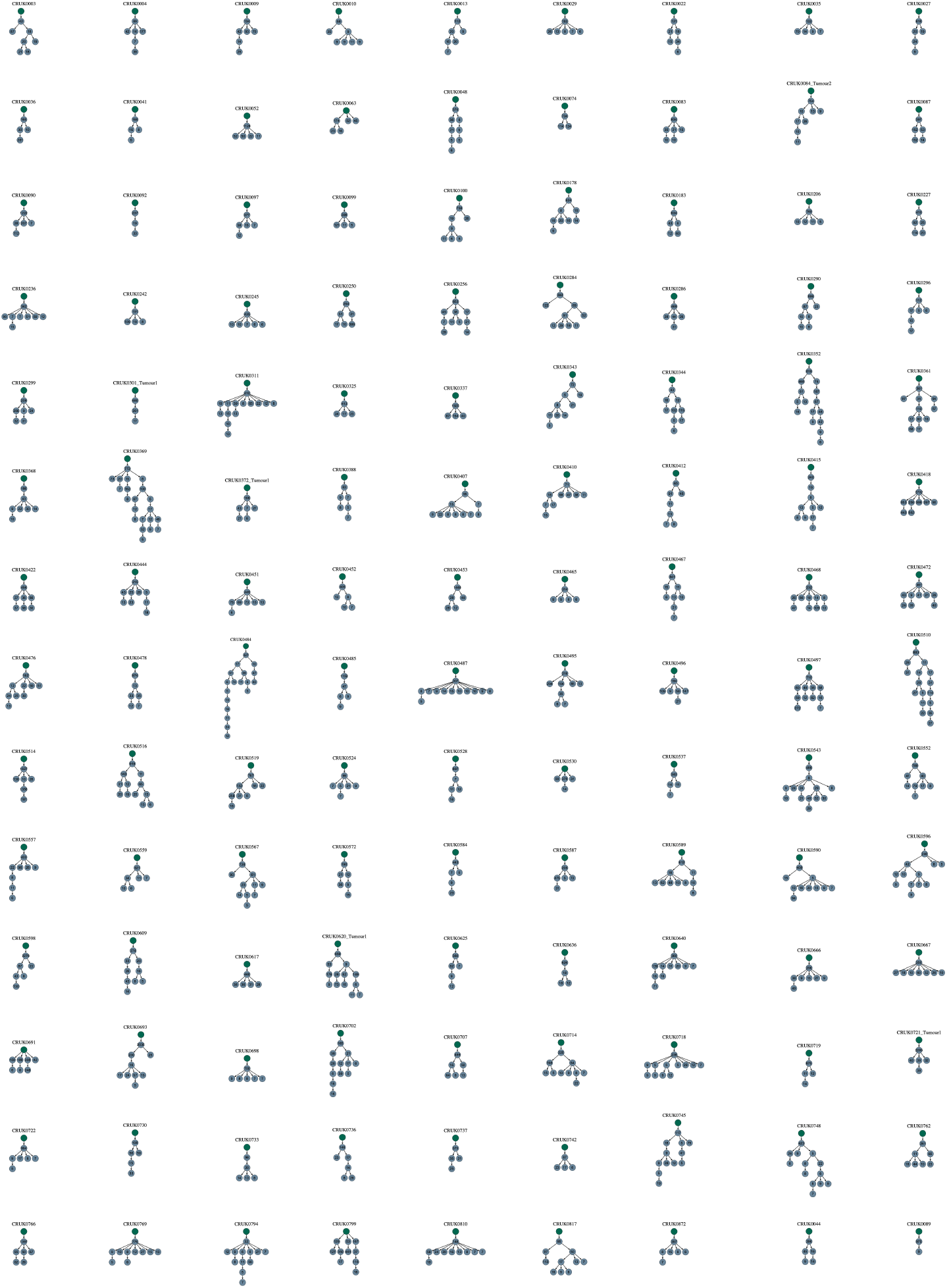
`omlta` of `CONIPHER` and `PairTree` inferred trees for each case in the NSCLC cohort. Nodes are labeled by the number of mutations in the cluster.

**Table S1:** Timing of first metastatic seeding event in each case of the NSCLC cohort, indicated by the CONIPHER and PairTree inferred trees, as well as their omlta.

| Case | CONIPHER | PairTree | omlta |
| --- | --- | --- | --- |
| CRUK0003 | late | early | early |
| CRUK0004 | late | late | late |
| CRUK0009 | early | late | late |
| CRUK0010 | late | early | late |
| CRUK0013 | late | late | late |
| CRUK0022 | late | late | late |
| CRUK0027 | late | late | late |
| CRUK0029 | early | early | late |
| CRUK0035 | early | early | late |
| CRUK0036 | late | early | late |
| CRUK0041 | late | late | late |
| CRUK0044 | late | late | late |
| CRUK0048 | late | late | late |
| CRUK0052 | late | early | late |
| CRUK0063 | late | early | early |
| CRUK0074 | late | late | late |
| CRUK0083 | late | early | late |
| CRUK0084_Tumour2 | late | early | late |
| CRUK0087 | early | early | late |
| CRUK0089 | late | late | late |
| CRUK0090 | early | early | late |
| CRUK0092 | late | late | late |
| CRUK0097 | late | early | late |
| CRUK0099 | late | late | late |
| CRUK0100 | late | late | late |
| CRUK0178 | late | early | early |
| CRUK0183 | late | early | late |
| CRUK0206 | late | late | late |
| CRUK0227 | early | late | late |
| CRUK0236 | late | late | late |
| CRUK0242 | late | early | late |
| CRUK0245 | late | late | late |
| CRUK0250 | early | early | early |
| CRUK0256 | late | early | early |
| CRUK0284 | late | early | late |
| CRUK0286 | late | early | late |
| CRUK0290 | early | early | early |
| CRUK0296 | late | early | late |
| CRUK0299 | late | early | late |
| CRUK0301_Tumour1 | late | late | late |
| CRUK0311 | late | early | late |
| CRUK0325 | early | early | late |
| CRUK0337 | early | late | late |
| CRUK0343 | late | early | early |
| CRUK0344 | late | early | late |
| CRUK0352 | late | early | late |
| CRUK0361 | late | late | late |
| CRUK0368 | late | early | late |
| CRUK0369 | late | early | early |
| CRUK0372_Tumour1 | late | late | late |
| CRUK0388 | late | NA | NA |
| CRUK0407 | late | late | late |
| CRUK0410 | late | early | late |
| CRUK0412 | late | early | early |
| CRUK0415 | late | early | early |
| CRUK0418 | late | early | late |
| CRUK0422 | early | early | late |
| CRUK0444 | late | early | late |
| CRUK0451 | early | early | late |
| CRUK0452 | late | late | late |
| CRUK0453 | early | early | early |
| CRUK0465 | early | early | late |
| CRUK0467 | early | early | early |
| CRUK0468 | late | early | late |
| CRUK0472 | early | early | late |
| CRUK0476 | late | late | late |
| CRUK0478 | late | NA | NA |
| CRUK0484 | late | early | late |
| CRUK0485 | early | early | late |
| CRUK0487 | early | early | late |
| CRUK0495 | late | early | early |
| CRUK0496 | late | early | late |
| CRUK0497 | late | early | late |
| CRUK0510 | late | early | late |
| CRUK0514 | late | late | late |
| CRUK0516 | late | early | early |
| CRUK0519 | late | early | late |
| CRUK0524 | late | early | late |
| CRUK0528 | late | early | late |
| CRUK0530 | early | early | late |
| CRUK0537 | late | early | late |
| CRUK0543 | late | early | late |
| CRUK0552 | late | early | late |
| CRUK0557 | late | early | late |
| CRUK0559 | early | early | early |
| CRUK0567 | early | early | early |
| CRUK0572 | late | early | late |
| CRUK0584 | late | early | late |
| CRUK0587 | early | early | late |
| CRUK0589 | late | late | late |
| CRUK0590 | late | early | late |
| CRUK0596 | late | early | early |
| CRUK0598 | late | early | late |
| CRUK0609 | early | early | early |
| CRUK0617 | late | late | late |
| CRUK0620_Tumour1 | late | early | late |
| CRUK0625 | early | early | late |
| CRUK0636 | early | late | late |
| CRUK0640 | late | late | late |
| CRUK0666 | late | early | late |
| CRUK0667 | late | late | late |
| CRUK0691 | early | early | late |
| CRUK0693 | early | early | early |
| CRUK0698 | late | early | late |
| CRUK0702 | late | early | late |
| CRUK0707 | late | early | late |
| CRUK0714 | early | early | early |
| CRUK0718 | late | early | early |
| CRUK0719 | late | late | late |
| CRUK0721_Tumour1 | late | late | late |
| CRUK0722 | late | late | late |
| CRUK0730 | late | late | late |
| CRUK0733 | late | early | late |
| CRUK0736 | late | early | late |
| CRUK0737 | early | late | late |
| CRUK0742 | early | late | late |
| CRUK0745 | late | early | late |
| CRUK0748 | late | early | late |
| CRUK0762 | late | early | late |
| CRUK0766 | late | late | late |
| CRUK0769 | late | late | late |
| CRUK0794 | late | early | late |
| CRUK0799 | late | early | late |
| CRUK0810 | early | early | late |
| CRUK0817 | early | early | early |
| CRUK0872 | late | late | late |

**Figure S5:**
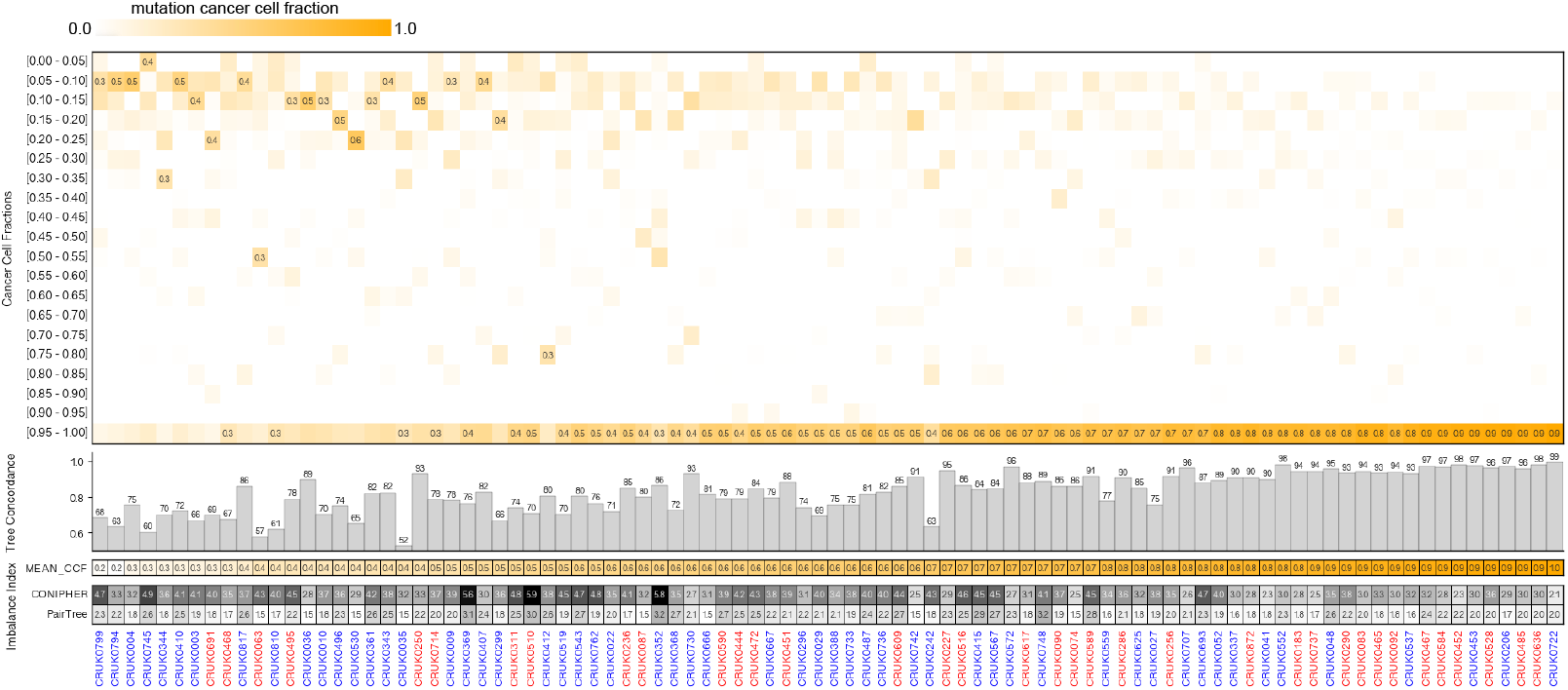
Each case of LUAD (colored blue) and LUSC (colored red) tumors from the NSCLC cohort, sorted by the mean CCF of mutations harbored. The top panel depicts the distribution of CCF across mutations harbored by each tumor. The next panel (gray bars) depicts the proportion of mutations that are preserved in the `omlta` of the `PairTree` and `CONIPHER` inferred trees for each case. The lowest panel depicts the average depth of a node in each tree inferred by each tool. As can be seen, the proportion of LUSC cases increases with increasing mean CCF. Similarly, the proportion of mutations that are preserved in `omlta` increases with increasing mean CCF. As a result the trees inferred for LUSC cases tend to be more robust to the choice of the tree inference method, and thus could be considered to be more reliable.

**Figure S6:**
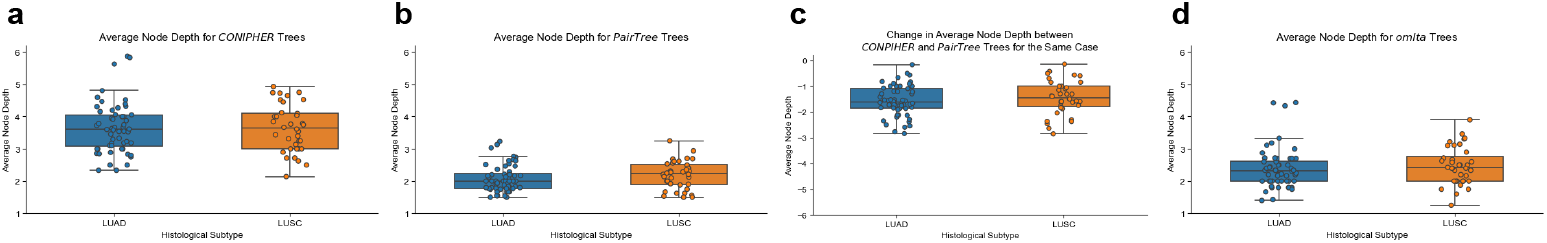
LUAD and LUSC trees inferred by the use of **a** `CONIPHER` and **b** `PairTree` do not demonstrate significant differences in tree imbalance (measured as the average node depth). **c**. Changes in tree imbalance for the same case between `CONIPHER` and `PairTree` trees are not statistically different between LUAD and LUSC. **d**. The `omlta` between `CONIPHER` and `PairTree` trees do not exhibit significant differences in tree imbalance across subtypes.

### A.5 Obtaining clonal trees from single-cell RNA sequencing data

This section includes more information on how we inferred trees on single-cell data.

#### A.5.1 Clustering cells in the genotype matrix

Variations in expression, allelic dropouts, and sequencing and/or genotyping errors are characteristic of single-cell RNA sequencing data (68). The high levels of sparsity and noise make it typically challenging to reliably disambiguate the true clonal structure of the tumor (60; 69; 19). We addressed this challenge by performing pre-clustering of cells as a step to building a robust tumor phylogeny. Since all of the tumors are derived from a single parental cancer cell line, we clustered cells from both control- and treated-mice together. Before building trees, we pre-clustered cells into mutationally homogeneous groups to mitigate the sparsity (which is a consequence of low sequencing coverage) in these datasets. The cells from both control and treated tumors were clustered together. In a later step, cells in each cluster were separated based on the treatment group and the respective trees were built independently.

The main input consists of genotype calls for each mutational locus in each cell in the form of a genotype matrix. A cell is represented as a ternary row vector with values corresponding to that cell’s mutational status at a genomic locus (‘0’ reference allele, ‘1’ mutated allele, and ‘NA’ missing data). We defined a pairwise similarity measure between two cells as the normalized Hamming distance computed by excluding sites that are ‘NA’s in either cell and normalizing by dividing the unnormalized Hamming distance number of informative cells. Through complete-link clustering, we established a hierarchy among cells. This hierarchy was subsequently partitioned into disjoint clusters, which is done by finding the minimum cophenetic distance (70) to merge any two clusters such that no more than *S* clusters form. In general, we chose a large *S* to avoid under-clustering, as a small number of clusters invariably limits the granularity to which tumor evolution can be resolved by the corresponding tree. For each cluster, we set its mutational profile by establishing a consensus for each mutational locus based on the combined read coverage over all cells in the cluster. We removed clusters that contained fewer than *c* (= “Min. cells per cluster” in Table 1) cells. Finally, we separated each cluster according to each dataset such that we build their respective trees independently. For each cluster, we defined its mutational profile by establishing a consensus for each mutational locus based on the combined read coverage over all cells in the cluster.

#### A.5.2 Tree inference

In this work, we inferred some clonal trees of tumor progression using `ScisTree` (60) which takes as input a genotype probability matrix where instead of the mutational state (e.g., 0/1/NA), each entry represents the probability of the mutation being present in the cell. These probabilities were parameterized by the noise rates related to sequencing; we empirically chose false positive and false negative rates consistent with the input, aiming to minimize the discrepancy between input and solution-implied values. In instances of missing entries in the genotype matrix, we simply set the probability to 0.5. `ScisTree` performs a heuristic, iterative search in the space of nearest neighbor interchange trees for the maximum likelihood, infinite-sites satisfying tree. The resultant conflict-free genotype matrix corresponds to a clonal tree labeled on the set of mutations. See Tables 1 and 2 for the `omltd` measures between inferred trees for each setting of the total number of clusters and the minimum number of cells allowed in a cluster, and see Figure 8 for the trees built from each sequencing dataset and their `omlta` alignment when the maximum number of the clusters, *S*, is set to 18.

### A.6 Comparison of omltd to previously defined measures of clonal tree similarity

Several measures for comparing clonal trees of tumor progression have been proposed recently (71; 22; 16). We have benchmarked `omltd` against six of these measures on simulated data generated with various parameter settings. Before we present our experimental results we discuss the strengths and weaknesses of these measures.

#### (i) Ancestor-Descendant accuracy and Different-Lineage accuracy

Perhaps the two most predominantly used measures of similarity between clonal trees are (a) Ancestor–Descendant accuracy, the proportion of pairs of mutations which have the same ancestor-descendant relationship in both input trees, among all pairs of mutations which have an ancestor-descendant relationship in either tree, and (b) Different-Lineage accuracy, the proportion of mutation pairs which occur at different nodes and do not have an ancestor-descendant relationship (i.e., are in different lineages) in both input trees, among all pairs of mutations which are on different lineages in either tree.When comparing an inferred tree with a known ground truth tree (e.g., in simulations), it is possible to define these relationships as a proportion of pairs of mutations with an ancestor-descendant or different-lineage relationship in the ground truth tree that have the same relationship in the inferred tree. These measures are considered to be complementary to each other and are typically used together to assess similarity between clonal trees, e.g., in (18; 72; 17). While these measures are easy to compute, some of their limitations have been previously described on a number of examples (16; 22). Nevertheless, because of their popularity, we have used these two measures for benchmarking in comparison to `omltd`. There are other related measures, such as Clustering Accuracy (18) and Co-Clustering Accuracy (73) for quantifying placement differences between mutation pairs originating at the same clone in one or both of the input trees. These measures typically have similar limitations to Ancestor-Descendant and Different-Lineage accuracy measures and are not commonly used.

#### (ii) Common Ancestor Set (CASet) and Distinctly Inherited Set Comparison (DISC)

A more recently introduced pair of complementary measures to account for structure of mutation inheritance are CASet and DISC distances (22), which collectively aim to address some of the shortcomings of the Ancestor-Descendant and Different-Lineage measures. (1) CASet compares two trees based on the ancestor sets of mutation labels in each tree. Informally, CASet considers each pair of mutations *i* and *j* and for each tree, considers the complete set of mutations that label *common* ancestors of the nodes labeled by *i* and *j*. Specifically, CASet is the difference (Jaccard distance) between the set of common ancestral mutations of *i* and *j* in the first tree, and that of the second tree, averaged across all mutation pairs *i, j*. Essentially, the CASet distance increases proportionally to the number of differences found between the ancestor sets of pairs of mutations in the two input trees. One observation about CASet is that even a single mutation can cause a drastic change in its distance calculation. The more descendants a node has, the more impact its labels have on pairs of mutations in the tree. This functionality of CASet distance was intended to reflect the potential increased significance of mutations that occur in the cell lineage earlier since they (typically) are present in a larger fraction of cells. It is easy to find many examples, however, where a single mutation swap in a tree can cause the distance between the original tree and the new tree to jump from 0 to nearly 1, the maximum distance value. Figure S7a provides one such example in which the root label is swapped with a leaf label, leading to the CASet distance to be larger than .9 even though the topology and labeling matches entirely otherwise. In comparison to CASet, one of the most noticeable benefits of tree edit distance is that it has a very low variance across different pairs of trees that have minimal differences in topologies and node labeling, which we later show experimentally. (2) Similar to Different-Lineage measure complementing the Ancestor-Descendant measure, DISC complements CASet in the sense that for a given pair of mutations *i* and *j*, it considers the complete set of mutations that label *distinct* ancestors of the nodes labeled by *i* and *j*. Specifically, DISC is the difference (Jaccard distance) between the set of distinct ancestral mutations of *i* and *j* in the first tree, and that of the second tree, averaged across all mutation pairs *i, j*. We have benchmarked both CASet and DISC against `omltd` and other measures we considered on simulated data.

**Figure S7:**
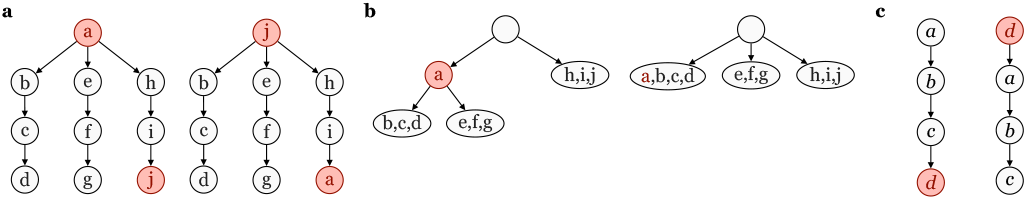
Example pairs of trees whose minor differences are better captured by `omltd` than alternative measures. **a**. (vs CASet) The two trees have very similar topologies and only differ by their placement of mutations *a* and *j*, which are swapped between the trees. However they have a CASet distance of 0.94 out of 1.0. In contrast their `omltd` (normalized by the maximum possible `omltd` between trees with 10 mutations) is only 0.2. **b**. (vs MLTD) The two trees only differ by the placement of mutation *a* as being ancestral to mutations *b, c, d* and *e, f, g* (through the addition of a new parental node) in the left tree as compared to the right tree. MLTD (normalized by the maximum possible MLTD between trees with 10 mutations) between these trees is 0.3, since either the entire set of mutations *d, c, b* or alternatively *g, f, e* need to be deleted (in this order) to make the trees isomorphic - subject to empty leaf deletions and node expansions. On the other hand the `omltd` (normalized by the maximum possible `omltd` between trees with 10 mutations) between these trees is only 0.1, a third of the MLTD, since the deletion of the node containing mutation *a* in the left tree makes them isomorphic. **c**. (vs Bourque Distance) The two trees differ by where mutation *d* is placed–in the left tree as the leaf and in the right tree as the root label. The pairwise Bourque Distance (normalized by the maximum Bourque Distance between trees with 8 mutations) is 1.0 since all edges in these trees need to be contracted. In contrast, `omltd` (again normalized by the maximum `omltd` between trees with 8 mutations) is only 0.25, reflecting that the deletion of mutation *d* from both trees makes them isomorphic.

#### (iii) Multi-Label Tree Dissimilarity (MLTD)

MLTD was a limited variant of tree edit distance proposed by (16) intended to provide a polynomial-time computable similarity measure for clonal trees. MLTD is one of the first specialized measures designed for comparing clonal trees and identifying their consensus tree. The main limitation of MLTD, as emphasized by the authors in their work, is that only leaf deletions are allowed rather than general node deletions. The benefit of this requirement is that the problem is no longer NP-hard unlike the original unordered tree edit distance. Computability aside, `omltd` offers a conceptual improvement to MLTD as a measure since by only allowing leaf deletions, it is possible that MLTD incurs a huge penalty for a small topological difference high up in the tree. Figure S7b provides an example in which only a single change in topology leads to an MLTD score triple that of the tree edit distance.

#### (iv) Bourque distance

The Bourque distance is a recent generalization for clonal tree comparison of the classic Robinson-Foulds tree similarity measure originally proposed for phylogenetic tree analysis. For a pair of rooted input trees, the Robinson-Foulds measure counts the symmetric distance between each tree’s set of two-part edge-induced partitions. For each edge (*u, v*) of a tree, the first part of its induced two-part partition is the label set of *v* and all descendants of *v*, and the second part is the remaining set of labels of the tree. Robinson-Foulds seeks to count how many such partitions are unique to each tree and reports the count as their similarity (23). As can be seen from its definition, both label placement and tree topology will affect the Robinson-Foulds metric. Robinson-Foulds cannot be used to measure clonal tree similarity primarily due to the fact that two clonal trees being compared may not have the same label set; according to its definition, the Robinson-Foulds score would be maximized by the appearance of even a single label found in one tree but not the other.

Bourque distance attempts to address the issue of Robinson-Foulds with a natural solution: only compute the bipartitions on the shared label set between two given input trees. The Bourque distance is an efficiently computable distance measure and serves as a useful modernization of a classic metric. However, similar to some of the other distance measures discussed above, we note that Bourque distance can be very sensitive to even a single mutation changing locations (see Fig. S7c).

**Figure S8:**
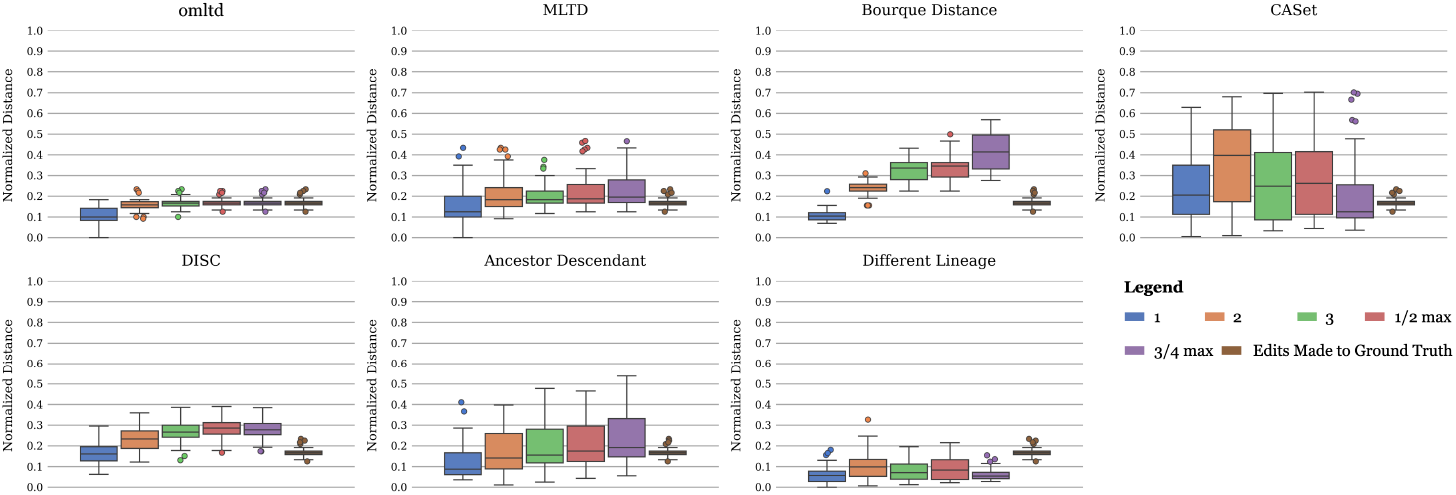
Each plot depicts the distribution of a particular measure of tree similarity across 250 simulated ground truth trees and their altered versions: each of the the leftmost 5 boxes in a plot corresponds to a particular setting of *d*, demonstrating the distribution of the measure across 5 distinct tree topologies, each with 10 different mutational assignments (blue for *d* = 1, orange for *d* = 2, green for *d* = 3, red for half the maximum possible distance in the tree, and purple for three quarters of the maximum possible distance). The rightmost box in each plot (in brown) corresponds to the true number of moved mutations between the ground truth tree and the altered tree.

### A.7 Comparisons of omltd with existing measures on simulated data

We employed the simulator introduced in a previous work (73), to generate 5 distinct tree topologies for tumor progression, each with 30 nodes; in each such tree, the root node represent the normal cells. For each of the simulated tree topologies *T*, we assigned 120 mutations to its non-root nodes. To ensure that each non-root node represents a genetically distinct population of cells, we first assigned one randomly chosen mutation to each such node (there are 30 ™1 = 29 such nodes/mutations); we then assigned each of the remaining (120 ™29 = 91) mutations to these nodes randomly. This forms the ground truth tree *T* . Next, for each node *v* in tree *T* we computed distances from *v* to all other nodes of *T* ; we denote by *d*_*v*_ the largest of these distances, and by *N*_*d*_(*v*) the set of all non-root nodes that are at distance exactly *d* from node *v*. Then, for each setting of *d* (which includes three fixed values, 1, 2, 3, as well as ⌊*d*_*v*_*/*2⌋ and ⌊3*d*_*v*_*/*4⌋, which are specific to node *v*), we altered *T* as follows. We randomly chose a set of 5 distinct non-root nodes in *T*, namely *V* = {*v*_1_, *v*_2_, …, *v*_5_} and for each *v*_*i*_ we randomly selected a distinct non-root node 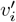 from the set *N*_*d*_(*v*_*i*_) which is not in *V* . Next, we moved all mutations in each *v*_*i*_ to 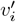 to obtain the altered version of *T*, namely *T* ^*′*^. While these two trees are topologically isomorphic, *T* differs from *T* ^*′*^ with respect to the described mutation assignments. For each setting of *d*, we repeated this entire process 10 times, thus obtaining 5 (tree topologies) × 5 (settings of *d*) × 10 (repeats) = 250 simulated instances.

For each simulated instance, comprised of the ground truth tree *T* and its altered version *T* ^*′*^, we computed their `omltd`, as well as CASet, DISC, MLTD, Bourque, Ancestor-Descendant, and Different-Lineage measures. Figure S8 depicts the results of these comparisons. In each of the seven plots, we display the distribution of the corresponding measure between *T* and *T* ^*′*^ for each of the 5 settings of *d*. In each plot, the rightmost box (in brown) corresponds to the distribution of the true number of mutations moved to obtain *T* ^*′*^ from *T* (normalized by the total number of mutations = 120) in the 250 simulated instances; as such, it depicts the distribution of the true distances between *T* and *T* ^*′*^ (thus the rightmost box depicts the same distribution in each of the seven plots). Among the seven measures tested, `omltd` matches the true number of moved mutations the best. Furthermore, it has the lowest variance and thus the highest reliability among the seven measures we tested across the simulated instances for each setting of *d* . Finally, as desired, `omltd` is not much impacted by the setting of *d*. However, especially when *d* = 1, the value of `omltd` may turn out to be lower than the true number of moved mutations; this is because the optimal solution as defined by `omltd` may be lower than the ground truth especially when *T* ^*′*^ is obtained from *T* by moving mutations between adjacent nodes (i.e., when *d* = 1).

After `omltd`, the tightest distributions are observed for DISC and Different-Lineage measures. Among these two, Different-Lineage values are typically much smaller than the true number of moved mutations, but with several outliers having much higher values. Similar to `omltd`, Different-Lineage is not impacted by the setting of *d*. DISC on the other hand is impacted by the setting of *d* and thus appears to be sensitive to where the mutations are moved. From the tightness of distribution point of view, Bourque distance and MLTD come next. For Bourque distance, not only the mean distance values but also their variances increase with increasing setting of *d*. This trend can be better understood through the toy example from our earlier discussion, in which moving a mutation to a farther node increase the distance. Thus, the Bourque distance would be useful only when mutational placements in an inferred tree differ from the ground truth locally. MLTD is relatively stable across different settings of *d* but its value typically higher than the true number of mutation moves and is not as tightly distributed as `omltd`. The final two measures, namely, Ancestor-Descendant, and CASet have a large ranges of values. While Ancestor-Descendant measure generally increases with the increasing setting of *d*, no such trend can be observed for CASet.

**Figure S9:**
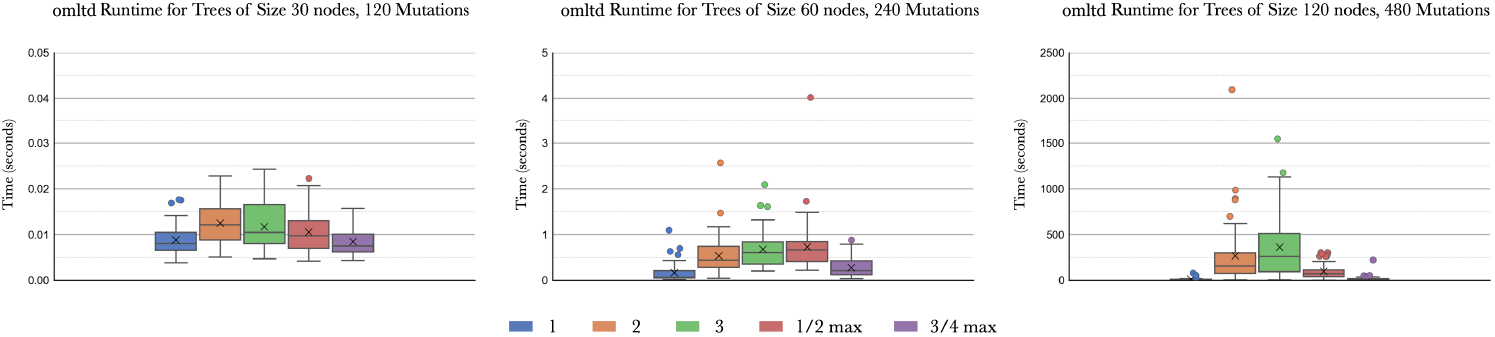
Each plot depicts the runtime distribution of our `omltd` algorithm on 50 simulations (on 5 distinct tree topologies, each with 10 distinct mutational assignments) obtained for each setting of *d* (blue for *d* = 1, orange for *d* = 2, green for *d* = 3, red for half the maximum possible distance in the tree, and purple for three quarters of the maximum possible distance). The number of nodes in the simulated trees as well as the total number of mutations double from the leftmost plot to the middle and from the middle to the rightmost plot.

### A.8 Runtime of omltd on simulated data with different sizes

We measured and compared the time taken to compute `omltd` on simulated datasets of increasing sizes. First, using the method described in Section A.7, we generated 250 ground truth trees (50 for each setting of *d*), each with 30 nodes and 120 mutations, and moved the mutations in 5 of these nodes to obtain the altered tree. Next we again used the same simulation method to generate another set of 250 ground truth trees, this time with 60 nodes and 240 mutations each, and moved the mutations in 10 of these nodes to obtain the altered tree. Finally we generated 250 more ground truth trees each with 120 nodes and 480 mutations, and moved the mutations in 20 of these nodes.

Figure S9 provides runtime results on these three simulated data sets across each setting of *d*. As expected, the runtimes increase substantially with the number of mutations moved (the mean number of mutations moved doubles from the left plot to the middle plot, and then again to the right plot) - since the runtime of our algorithm depends exponentially on the value of `omltd`. Interestingly the runtime also depends on the value of *d*, i.e., the distance between the original placement of mutations and the final placement. The runtime peaks at *d* = 3 but gets smaller as *d* changes in either direction. The impact of the value of *d* is substantial on the largest trees (rightmost plot) where it may change the runtime by several orders of magnitude. This is likely because (i) when the mutations are moved to more distant nodes, the number of low-cost edit sequences are substantially reduced, enabling `omltd` algorithm to explore fewer options, and (ii) when the mutations are moved to neighboring nodes, the search space is again limited.

